# Prelimbic-amygdala overexcitability mediates trait vulnerability in a novel mouse model of acute social defeat stress

**DOI:** 10.1101/2020.06.11.147231

**Authors:** Yael S. Grossman, Clementine Fillinger, Alessia Manganaro, George Voren, Rachel Waldman, Tiffany Zou, William Janssen, Paul Kenny, Dani Dumitriu

## Abstract

**BACKGROUND:** Depression is a debilitating neuropsychiatric disorder with 20% lifetime prevalence in the developed world but only approximately half of afflicted individuals respond to currently available therapies. While there is growing understanding of the neurobiological underpinnings of the depressed brain, much less is known about the preexisting circuitry leading to selective vulnerability versus resilience. Elucidating these networks could lead to novel preventative approaches.

**METHODS:** We developed a model of acute social defeat stress (ASDS) that allows classification of male mice into “susceptible” (socially avoidant) versus “resilient” (expressing control-level social approach) one hour after exposure to six minutes of social stress. Using circuit tracing and high-resolution confocal imaging, we explored differences in activation and dendritic spine density and morphology in the prelimbic to basolateral amygdala (PL→BLA) circuit in resilient versus susceptible mice. To test the functional relevance of identified structure/function differences to divergent behavioral responses, we used an intersectional chemogenetic approach to inhibit the PL→BLA circuit during or prior to ASDS.

**RESULTS:** Susceptible mice had greater PL→BLA recruitment during ASDS and activated PL→BLA neurons from susceptible mice had more and larger mushroom spines compared to resilient mice. Inhibition of the PL→BLA circuit led to a population shift towards resilience.

**CONCLUSION:** Preexisting PL→BLA structure/function differences mediate divergent behavioral responses to ASDS in male mice. These results support the PL→BLA circuit as a biomarker of trait vulnerability and potential target for prevention of stress-induced psychopathology.

## Introduction

Major depressive disorder (MDD) is the leading cause of years with disability worldwide[76] and roughly a third of affected individuals are resistant to therapeutic interventions[40]. Even treatment-responsive patients often require years of trial-and-error to identify an effective treatment. Several preexisting traits, features of cognitive development, and environmental factors have been implicated for conferring resilience or vulnerability to depression[15, 35, 39, 55, 64, 80, 82, 81]. This suggests that actions taken prior to symptomatic expression of MDD can have drastic effects on a patient’s prognosis. Identifying the vulnerable population could facilitate development of novel therapeutics, with the potential for alleviating[68], delaying[4], or preventing depression in at-risk individuals[54].

Chronic social defeat stress (CSDS) is a well-validated mouse model for depressive-like behaviors and one of few validated models for reproducible classification of two distinct stress-induced phenotypes[30, 49]. Following repeated exposure to novel aggressive mice, approximately two thirds of experimental mice exhibit susceptible behavior, defined as acquired social avoidance, while one third exhibit resilient behavior, defined as persistent social approach similar to control mice. Accumulating evidence points to vast differences in numerous neurocircuits in resilient versus susceptible mice following CSDS[14]. However, much less is known about the preexisting circuits that mediate downstream divergent stress-responses and circuit plasticity.

The prelimbic to basolateral amygdala (PL→BLA) circuit is necessary for the acquisition of fear memory[1, 45, 85], involved in anxiety-like behavior[1, 2, 66, 71, 88], and overactive in depressed patients[12, 20, 35, 53, 72, 84, 89]. Furthermore, PL neurons have stunted dendritic arbors after CSDS[14, 16, 73, 95]. Therefore, preexisting differences in the function and structure of the PL→BLA circuit are good candidates for investigating mechanisms underlying divergent stress-responses.

We developed a model of acute social defeat stress (ASDS) that allows for rapid classification of susceptible and resilient mice, enabling mechanistic dissection of pre-existing PL→BLA circuit structure/function differences and their contribution to divergent stress responses. Using tract-tracing in conjunction with the transcriptional activity marker cFos[19, 60], we show that mice susceptible to ASDS have greater proportion of activated PL→BLA neurons compared to resilient mice. Furthermore, activated PL→BLA neurons are morphologically different in susceptible versus resilient mice. Using an intersectional chemogenic approach, we then show both acute and chronic inhibition of this circuit leads to a population shift towards resilience. Together, these results implicate the PL→BLA circuit as a biomarker of trait vulnerability and a promising target for prevention of stress-induced maladaptive behavior.

## Methods

See Extended methods in Supplementary Materials for detailed methods.

### Animals

Experimental mice were 7-12 week old C57BL/6J male mice, group housed (4mice/cage), maintained on a 12 hour light/dark cycle, with *ad libitum* food and water. Retired breeder CD1 male mice that displayed aggression during screening were used in social defeat experiments and non-aggressive male CD1 mice were used in social interaction (SI) testing. All experiments were conducted in compliance with National Institutes of Health Guidelines for Care and Use of Experimental Animals approved by Institutional Animal Care and Use Committee at Icahn School of Medicine at Mount Sinai.

### Chronic social defeat stress (CSDS)

CSDS was performed as previously described[30, 49, 51]. Experimental mice were placed into cages of novel aggressive CD1 mice for 5 min daily for 10 consecutive days. For the remainder of each 24 hour period, intruder and resident aggressor remained co-housed with a clear perforated plexiglass partition prohibiting further aggression. Control mice interacted and were co-housed with novel non-aggressive con-specific mice daily. On day 10, experimental mice were group-housed with prior cage-mates, then tested on social interaction (SI) on day 11.

### Subthreshold social defeat stress (StSDS)

StSDS has been previously described as a submaximal stressor that does not lead to social avoidance and is used to assess increased susceptibility in response to various manipulations[30, 49, 52]. Experimental mice were placed into cages of three aggressive CD1 mice for 5 min each, with 15 min rest between aggressive sessions. Control mice were placed into cages of three novel non-aggressive conspecific mice. Experimental and control mice were then returned to group-housing with prior cage-mates and tested on SI 24 hours later.

### Acute social defeat stress (ASDS)

Various forms of acute social stress have been described[22, 63, 83], but a standardized ASDS protocol resulting in distributions of resilience and susceptibility similar to CSDS has not previously been validated to the best of our knowledge. Experimental mice were placed into the cages of three aggressive CD1 mice for 2 min each sequentially, without rest periods. Control mice were placed into cages of three non-aggressive conspecific mice for 2 min each sequentially. All mice were then singly-housed in a clean cage for 54 min, then tested on SI exactly 60 min following onset of ASDS.

### Behavioral assays

SI was tested identically for CSDS, StSDS and ASDS. Experimental mice were placed into an opaque Plexiglas open-field arena (42×42×42cm^3^) with a removable wiremesh enclosure placed against the middle of an inner wall of the arena. Mice first explored the arena for 150sec with “no target” present, then explored for 150sec with a novel non-aggressive CD1 “target” mouse in the enclosure. Video-tracking (Ethovision 3.0, Noldus Information Technology) was used to record movements. Total time spent in “interaction zone” (8cm area surrounding enclosure) and “corner zones” (9×9cm corners on opposite wall from enclosure) were quantified. Interaction and corner ratios were calculated by dividing time spent in respective zone during “target” present by time during “no target”.

For experiments validating ASDS, “resilience” was defined as SI ≥ 1 and “susceptibility” was defined as SI ratio<1 in accordance with the conventional classification for CSDS[49]. For experiments looking at PL→BLA activation and dendritic spine structure, we used a multidimensional classifier to enrich for behavioral homogeneity. The classifier and its rationale are described in detail in Extended Methods.

For sucrose preference testing, mice were individually housed to enable quantification of consumed fluids. Mice were first habituated to drinking water from two 50mL conicals with sipper stops for two days. The content of one conical was then replaced with 1% sucrose in water and fluid consumed from each conical was recorded at the end of a 24 hour period. In open field testing, mice explored an empty opaque Plexiglas arena (42×42×42cm^3^) for 5 min and time spent in center (10×10cm) was quantified using video-tracking. In elevated plus-maze testing, mice were placed in the center of a custom-built Plexiglass apparatus with two open arms and two enclosed arms 1m above floor level. Time spent in open arms during 5 min exploration was quantified using video-tracking.

### Viral-mediated tract tracing

Mice were injected with the retrograde virus AAV5-hSyn-eGFP (cat#AV-5-1696, University of Pennsylvania Vector Core) into the BLA unilaterally (counterbalanced, from Bregma: medio-lateral +/−3.4, anterio-posterior −1.1, dorso-ventral −5.0, angle 0°) under ketamine/xylazine anesthesia to label PL→BLA neurons. Mice were allowed to recover and fully express the virally-delivered GFP for minimum 18 days. Post-hoc histological confirmation of BLA targeting was performed on all brains and only those meeting both viral localization and behavioral classification criteria and were included.

### Chemogenetic inhibition of PL→BLA pathway

An intersectional chemogenetic approach was employed to specifically target the inhibitory Designer Receptor Exclusively Activated by Designer Drug (DREADD) hM4D(Gi) to PL→BLA neurons. AAV5-hSyn-Cre-GFP (cat#AV-5-PV1848, University of Pennsylvania Vector Core) was injected into BLA bilaterally and AAV8-hSyn-DIO-hM4D(Gi)-mCherry (cat#44362, Addgene) was injected into PL bilaterally (from Bregma: medio-lateral +/−0.8, anterio-posterior +2.3, dorso-ventral −2.3, angle 11°). Mice were allowed a minimum of three weeks to recover and fully express virally-delivered gene products.

For acute PL→BLA inactivation, Clozapine-N-oxide (CNO, 2 mg/kg, dissolved in saline) or saline was injected intraperitoneally (i.p.) 30 min prior to ASDS. Four mice were used for assessing the effect of CNO on PL and PL→BLA cFos and for immunohistochemical evaluation of fidelity and efficiency of the intersectional approach. Remaining mice were allowed one week of recovery, followed by StSDS in the absence of CNO. For chronic PL→BLA inactivation, CNO (0.25mg/ml) or vehicle (aspartame 0.005%) was provided in drinking water ad libitum for 10 days. On day 11, mice were subjected to ASDS in absence of CNO. Post-hoc histological confirmation of injections sites was performed on all brains and only those with adequate localization of all four injection sites were used in behavioral analyses of DREADD-positive mice. Mice that did not show any mCherry expression anywhere, i.e. failed intersectional chemogenetic approach, and were used as DREADD-negative virally-injected controls.

### Tissue collection and processing)

For tract-tracing and chemogenetic studies, mice were anesthetized with chloral hydrate (1.5g/kg) and sacrificed by transcardial perfusion with paraformaldehyde immediately after SI testing. Brains were post-fixed for 6 hours and sectioned at 150 *μ*m. Immunohistochemistry was optimized for antibody penetration in thick sections to allow visualization of entire neurons (see Extended Methods). Sections were serially stained with rabbit anti-cFos primary antibody (cat#sc-52, Santa Cruz) and Alexa Fluor-conjugated goat anti-rabbit secondary antibody (Alexa Fluor-568, cat#A-11011 for tract-tracing, or Alexa Fluor-647, cat#A-21244 for chemogenetic studies, Life Technology), followed by rabbit anti-GFP primary antibody (cat#A-11122, Life Technology) and Alexa Fluor-488-conjugated goat anti-rabbit secondary antibody (cat#A-11008, Life Technology). For all experiments, sections were incubated in 4’,6-diamidino-2-phenylindole (DAPI) for histological identification.

### Imaging and image analysis

A Zeiss LSM510 confocal microscope was used for all imaging. Exact settings are included in Extended Methods. For PL→BLA cFos quantification, low resolution (10X) sequential dual-channel 3-dimentional (3D) tile scans were acquired of the entire medial prefrontal cortex (mPFC). For dendritic spine imaging, neighboring PL→BLA cFos immunoreactive (cFos+) and cFos non-immunoreactive (cFos–) pyramidal neurons with somas in layers II-V spanning bregma 1.9-2.2 were targeted for high resolution (100x) 3D-stacks of apical and basal dendrites.

For chemogenetic experiments, triple/quadruple-channel 3D-stacks were acquired to image GFP, mCherry, and cFos, and sometimes DAPI. For histological confirmation of injection sites, low resolution (5X) images of sections spanning entire anterio-posterior extent of PL and BLA were inspected. For CNO effects on cFos expression and the fidelity and efficiency of the intersectional chemogenetic approach, higher resolution (25X) images were acquired.

Images acquired for cFos quantification and chemogenetic experiments were imported into Fiji[79], segmented into composite channels, smoothed via mean filtering, then top-hat filtered. Object identification was performed using Foci Picker 3D[21], saved as 3D 8-bit gray-scale tiffs, and exported to MatLab for 3D-colocalization (defined as >50% overlap) using custom code. Dendritic spines were quantified semi-automatically using NeuronStudio similarly to methods previously described[23].

### Statistical analyses

Percent activation of PL→BLA neurons and spine density were calculated in Excel. Statistical analysis was performed using MatLab2018b or GraphPad Prism8. One-way analyses of variance (ANOVAs) were used with group as the variable for comparison of percent activation and dendritic spine density. One-way and two way ANOVAs with Bonferroni post-hoc T-tests were used to compare group responses to ASDS and CSDS. Behavioral responses to ASDS in presence/absence of CNO were compared using two-way ANOVAs. Tukey post-hoc T-tests were used to detect differences between groups. Two-sample Kolmogorov-Smirnov (K-S) tests were used to test for significant differences in distributions of spine densities, spine head diameters, and spine length. In all statistical tests, threshold was set at p<0.05 and p-values were adjusted to correct for multiple comparisons, where necessary, using post-hoc Bonferroni correction.

## Results

### Development of standardized ASDS protocol for rapid identification of susceptibility and resilience

Our first objective was to develop and validate an acute stressor that identifies resilient and susceptible mice at a time-point when the protein for transcriptional activity marker cFos is maximally expressed[3, 69]. In our ASDS model, male intruder C57BL/6J mice are placed into the cages of three aggressive CD1 male resident mice for two minutes each, sequentially, and tested for social avoidance one hour after onset of stress (Figure 1). As a group, defeated mice become similarly socially avoidant following ASDS compared to CSDS (Supplemental Figure 1), but also display large individual variability in avoidant behavior (Figure 1C,D, Supplemental Figure S1C,D). The proportion of susceptible versus resilient mice (using conventional SI ≥ 1 to define resilience) is remarkably similar in ASDS versus CSDS, with 41 ± 5% (range 9-83%) resilience in ASDS and 42 ± 10% (range 28-93%) resilience in CSDS (Figure 1D, Supplemental Figure S1D). Resilient and susceptible mice also show similar patterns of interaction and corner zone durations (Figure 1E, Supplemental Figure S1E).

**Fig. 1.**
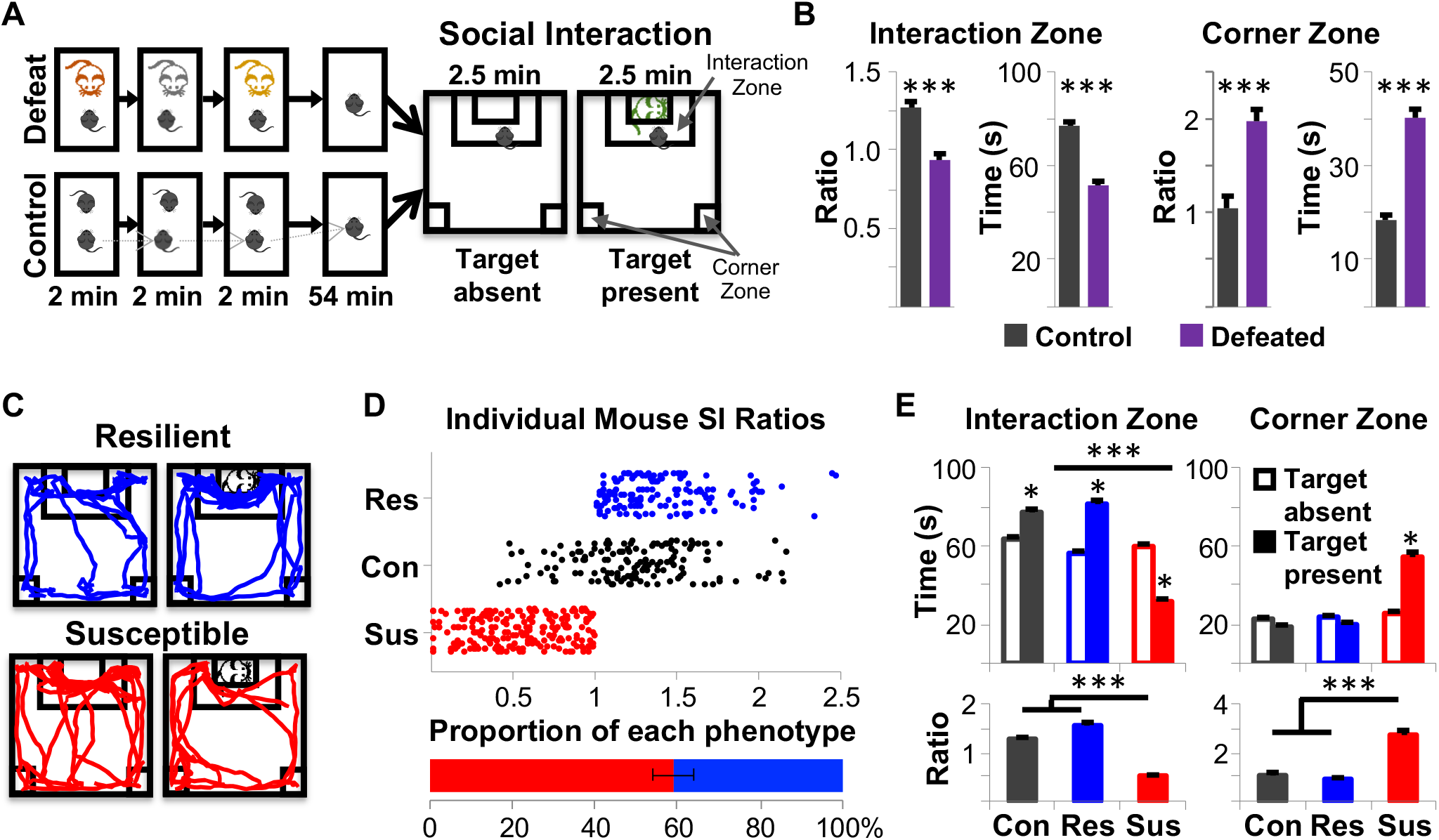
Validation of a novel model of Acute Social Defeat Stress (ASDS) in male mice. (A) Intruder C57 mice (black mouse in figure) are placed consecutively into home cages of three novel resident mice. For defeat, resident mice are retired breeder male CD1s that screened positive for aggression. For control, resident mice are age-matched non-aggressive male C57s. The intruder C57 spends two minutes in each of the resident home cages and aggression occurs during each encounter. The intruder mouse is then singly housed in a clean, empty cage for 54 minutes and immediately tested on social interaction (SI). For SI, the intruder C57 is first placed into an arena containing an empty enclosure and allowed to explore for 150 seconds. A non-aggressive CD1 “target” mouse is then placed in the enclosure and the intruder C57 is allowed to explore for another 150 sec. Movement is video-recorded and the time spent in interaction zone and corner zones is quantified. (B) Defeated intruder C57 mice spend significantly less absolute and ratio of time in interaction zone, and more absolute and ratio of time in the corner zone, when target present versus absent compared to controls (n=224 defeated, n=111 control, two-tailed unpaired t-test, p<0.0001 for all comparisons). (C) Susceptibility is defined as SI ratio<1. Resilience is defined as SI ratio ≥ 1. Representative movement of a susceptible and a resilient mouse in the arena when target is absent versus present, showing the typical social approach of a resilient mouse and the social avoidance of a susceptible mouse. (D) Distribution of SI ratios of individual control (Con, black), resilient (Res, blue), and susceptible (Sus, red) mice. Defeated mice are predominantly susceptible (59% ± 5%). (E) Absolute and ratio time spent interacting with target mouse and absolute and ratio time spent in corner zones shown following classification into susceptible versus resilient. Two-way ANOVAs were performed to compare interaction and corner times and one-way ANOVAs were performed to compare ratio scores. Bonferroni post-hoc tests were performed and all ANOVAs. Control and resilient mice spent significantly more time in the interaction zone when target was present than susceptible mice [two-way ANOVA, GroupXTarget interaction F(2,332)=607.7 p<0.0001, group main effect: F(2,332)=265.7 p<0.0001, target main effect: F(1,332)=11.03 p=0.001, post-hoc CvsR n.s., CvsS p<0.0001, RvsS p<0.0001], but not when target was absent, and higher SI ratios [one-way ANOVA, F(2,332)=5.932 p=0.0029, post-hoc CvsR n.s., CvsS p<0.0001, RvsS p<0.0001]. Conversely, susceptible mice spent more time in corner zones [two-way ANOVA, GroupXTarget interaction F(2,332)=10.9 p<0.0001, group main effect: F(2,332)=127.7 p<0.0001, target main effect: F(1,332)=48.68 p<0.0001, post-hoc CvsR n.s., CvsS p<0.0001, RvsS p<0.0001] and had lower corner time ratios [one-way ANOVA, F(2,332)=77.43 p<0.0001, post-hoc CvsR n.s., CvsS p<0.0001, RvsS p<0.0001] compared to susceptible mice. n.s.=not significant; C=control; R=resilient; S=susceptible; *p<0.05; ***P<0.001

### ASDS is not associated with pervasive depressive-like or anxiety-like behaviors but does prime for increased susceptibility to future social stress

CSDS is well-known to lead to pervasive depressive- and anxiety-like behaviors in subsets of mice(14). Therefore, we next asked if ASDS also leads to long-term maladaptive behaviors (Figure 2A). One week after ASDS, control and defeated mice did not differ on SI, sucrose preference, open field, elevated plus maze, or in body weight (Figure 2B).

**Fig. 2.**
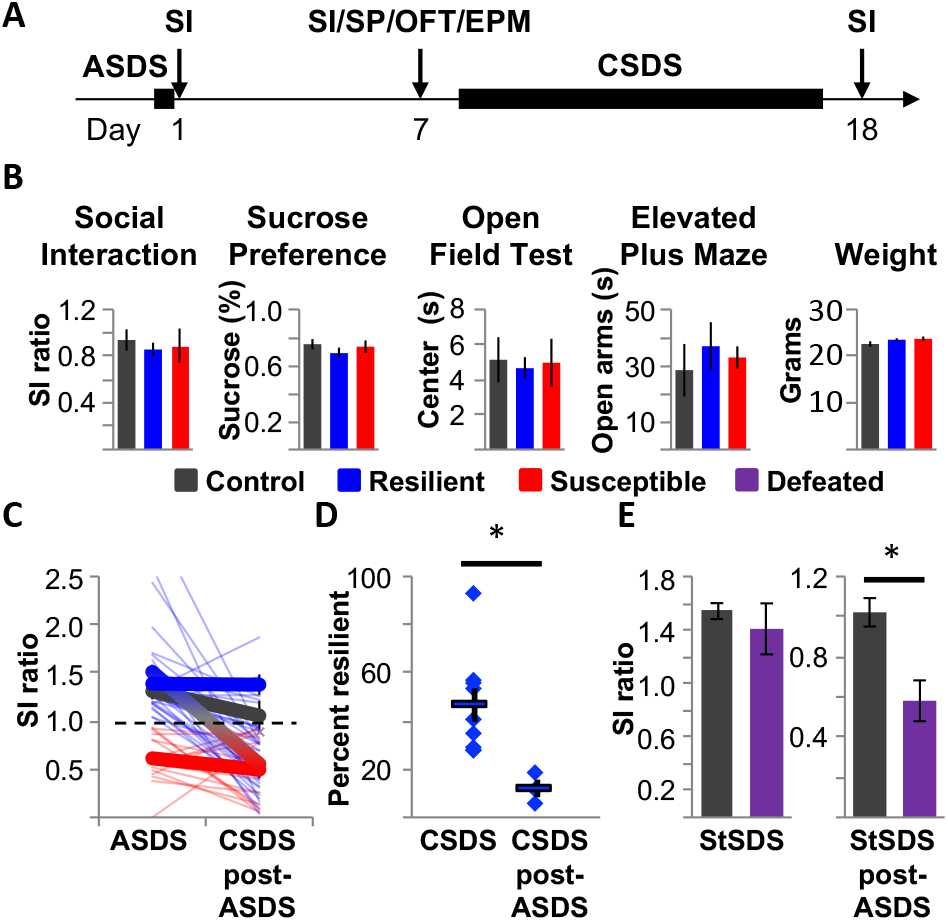
ASDS is not associated with pervasive depressive-like or anxiety-like behaviors but does prime for increased susceptibility to future social stress. (A) Experimental timeline. Mice first underwent ASDS and SI testing on Day 1. One week later, weights (control n=24, susceptible n=28, resilient n=44) were collected and different subsets of mice were tested on SI (control n=20, susceptible n=16, resilient n=34), sucrose preference (SP, control n=18, susceptible n=17, resilient n=35), open field test (OFT, control n=10, susceptible n=9, resilient n=22), and/or elevated plus maze (EPM, control n=6, susceptible n=10, resilient n=8). A subset of mice were then placed into chronic social defeat stress (CSDS). CSDS concluded on Day 17 and mice underwent SI on Day 18. (B) There was no significant difference between control, resilient, and susceptible mice in any of the tests performed on Day 7 [one-way ANOVA, SI F(2,67)=0.24 p=0.79; SP F(2,67)=0.67 p=0.51; OFT F(2,38)=0.071 p=0.93; EPM F(2,21)=0.27 p=0.77; Weight F(2,93)=1.56 p=0.22]. (C) All mice classified as susceptible after ASDS were also classified as susceptible after CSDS. Only 18% of mice that were classified as resilient after ASDS remained resilient after CSDS, with 82% becoming susceptible. (D) Percentage of CSDS-defeated mice classified as resilient when CSDS occurs post-ASDS (12±3.6%) is significantly lower than when CSDS is performed on naïve mice (42±10%, two-way unpaired T-test, p=0.021). (E) Naïve mice do not show social avoidance following StSDS (two-way unpaired T-test, p=0.53). In contrast, when StSDS occurs post-ASDS, defeated mice have significantly lower SI ratios (two-way unpaired T-test, p=0.038). *p<0.05

To assess if resilience to ASDS is predictive of resilience to CSDS, mice first underwent ASDS followed by CSDS after one-week recovery. While all mice classified as resilient following CSDS had been resilient to ASDS (Figure 2C), the comparison was limited by a significant increase in susceptibility (Figure 2D, Supplemental Figure 1D). “Super-resilient” mice, i.e. those remaining resilient after ASDS and CSDS, could not be predicted using any behavioral characteristics from SI following ASDS. However, resilience on initial ASDS did confer some protection regarding degree of social avoidance after CSDS (Supplemental Figure 2).

To evaluate if increased susceptibility represents ASDS-induced priming to future social stress, we tested mice that had undergone ASDS on a validated subthreshold paradigm (StSDS) that does not routinely induce susceptibility[13, 27]. As expected, StSDS did not lead to social avoidance in naïve animals. However, a significant decrease in SI ratio was seen in mice with a history of ASDS (Figure 2E).

### Susceptible mice have a greater proportion of PL→BLA neurons activated during ASDS

To assess if differential activation of the PL→BLA pathway, a circuit involved in fear memory formation[1, 10, 26, 45, 85], is associated with divergent stress-responses to ASDS, we employed viral-mediated tract-tracing (Figure 3A). Unilateral counterbalanced injections of a retrograde virus were used to quantify contra-versus ipsilateral PL→BLA activation. Viral injections did not affect ASDS-induced behavioral responses (Supplemental Figure 3). Prefrontal sections from control, susceptible and resilient mice with histologically confirmed BLA injections (Figure 3B) were double-stained against GFP and cFos (Figure 3C).

**Fig. 3.**
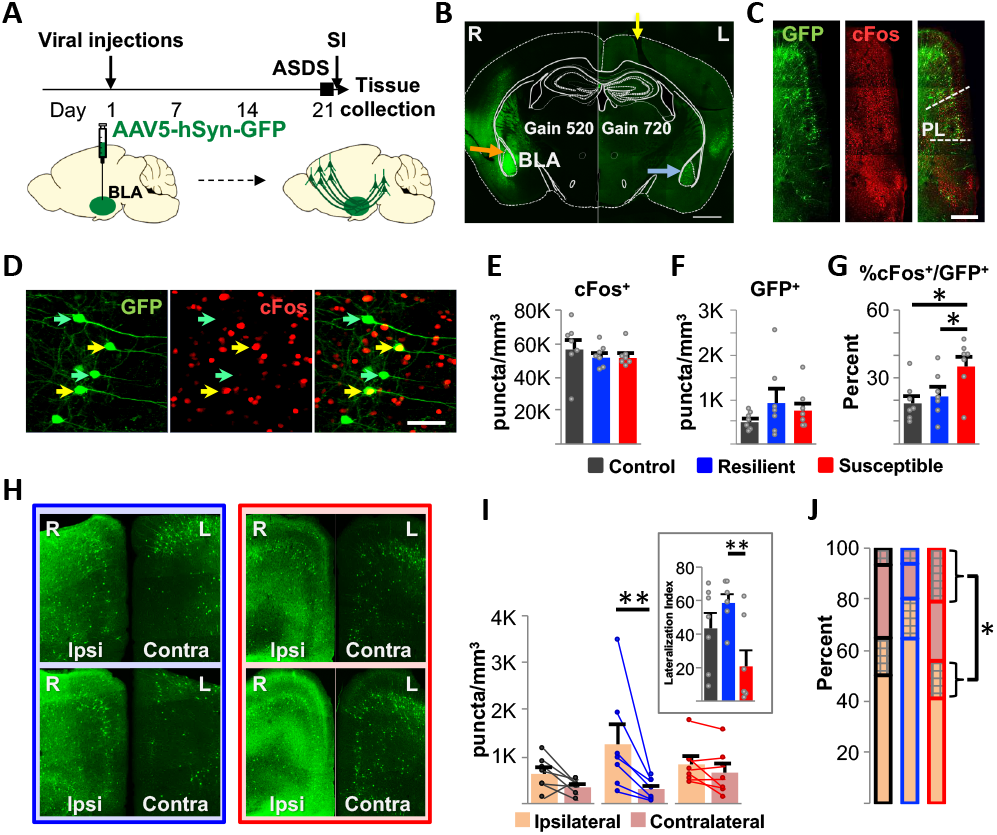
Susceptible mice have greater proportion of basolateral amygdala-projecting prelimbic (PL→BLA) neurons activated during ASDS. (A) Schematic of experiment design and timeline. A retrograde GFP-expressing virus was injected unilaterally in the BLA on Day 1. Twenty-one days later, mice underwent ASDS and SI, followed by immediate sacrifice and tissue collection. (B) Representative image of BLA injection. Outline overlay was taken from The Mouse Brain, 2nd Ed by George Paxinos and Keith Franklin (scale bar 1mm, plate 42, Bregma −1.34). Hemispheres were marked with a “nick” in the left hemisphere (yellow arrow, showing cut in retrosplenial cortex). Post-hoc histological confirmation of injection site was performed on all brains and inclusion criteria were defined as dense expression of GFP in injected BLA (orange arrow, hemisphere imaged at a lower gain of 520) and in contralateral BLA (blue arrow, hemisphere imaged at higher gain of 720 to show BLA → BLA projecting neurons). (C) Representative image (scale bar 0.5mm) of anti-GFP and anti-cFos stained medial prefrontal cortex (mPFC) with demarcated PL (Paxinos atlas, plate 14, bregma 1.98). (D) Representative higher resolution image (scale bar 200μm) of anti-GFP and anti-cFos staining in PL. Images were analyzed in 3D and PL→BLA neurons (as evidenced by anti-GFP staining) were classified as cFos positive (cFos+, yellow arrows) or cFos negative (cFos–, green arrows). (E) No significant difference in number of PL cFos puncta was observed between control, resilient, and susceptible mice [one-way ANOVA F(2,18)=1.07, p=0.36]. (F) No significant difference in number of GFP neurons was observed between control, resilient, and susceptible mice [one-way ANOVA F(2,18)=0.54 p=0.59]. (G) Susceptible mice had significantly higher proportion cFos+ GFP-expressing neurons than resilient and control mice [one-way ANOVA F(2,18)=5.01 p=0.019, post-hoc Bonferroni test: C vs S 0.012, R vs S 0.042, C vs R 0.56]. (H) Representative images of ipsilateral and contralateral GFP expression in mPFC of two resilient mice (blue outline) and two susceptible mice (red outline). (I) Resilient mice had significantly more GFP-expressing neurons in the ipsilateral compared to the contralateral hemisphere, whereas control and susceptible mice had comparable numbers in the two hemispheres [repeated measures two-way ANOVA with post-hoc Bonferroni correction, F(2,18)= 7.981, p=0.056, C p=0.68, R p=0.0017, S p>0.9999]. Inset: Lateralization was calculated as GFP puncta density in the ipsilateral hemisphere minus GFP puncta density in the contralateral hemisphere divided by total number of GFP puncta in both hemispheres. Resilient mice had significantly higher lateralization than susceptible mice [one-way ANOVA, F(2,18)=5.54 p=0.013, post-hoc Bonferroni test: RvsS p=0.004]. (J) Percentage of GFP-expressing neurons that co-express cFos was calculated in ipsilateral versus contralateral hemispheres in in each group. Percent co-expression did not differ between hemispheres in control (mean±SEM= 18±6% versus 22±5%, in contra-versus ipsi-lateral neurons respectively, two-tailed paired T-test p=0.70) or resilient (mean±SEM= 31±12% versus 20±7%, in contra-versus ipsi-lateral neurons respectively, two-tailed paired T-test p=0.10) mice. However, susceptible mice had significantly greater percent co-expression in the contralateral compared to the ipsilateral hemisphere (mean±SEM= 47±18% versus 26±10%, in contra- versus ipsi-lateral neurons respectively, two-tailed paired T-test p=0.01). C=control; R=resilient; S=Susceptible; *p<0.05; **P<0.01

Somewhat surprisingly, we found no difference in density of cFos puncta in the PL between any of the groups, suggesting physical aggression does not activate the PL above activation resulting from exposure to a novel social experience (Figure 3D-E). There was also no difference in PL density of GFP-labeled neurons between groups (Figure 3F). Consistent with our hypothesis, susceptible, but not resilient mice, had significantly higher proportion of GFP-labeled neurons double-stained for cFos compared to control mice (Figure 3G). This difference was specific to PL, with no differences in proportion of activated BLA-projection neurons in neighboring anterior cingulate (AC) or infralimbic (IL) mPFC subregions (Supplemental Figure 4A).

While overall number of PL→BLA neurons did not differ, we found an intriguing lateralization of projection neurons in resilient but not susceptible mice (Figure 3H, I).This pattern was most evident in PL, but similar trends were observed in AC and IL (Supplemental Figure 4B). Furthermore, we found an aberrantly high activation of contralateral PL→BLA neurons in susceptible mice (Figure 3J). Ipsi-/contra-lateral PL→BLA activation differences were not mediated by overall differences in PL ipsi-/contra-lateral activation or by left/right hemispheric activation of either PL or PL→BLA neurons (Supplemental Figure 5).

### Dendritic spine differences suggest higher excitatory drive in PL→BLA neurons of susceptible mice

We next investigated differences in dendritic spines, the primary sites of excitatory synaptic inputs to cortical neurons[8, 34, 44], as a potential structural mechanism for observed higher PL→BLA activation in susceptible mice. Apical and basal dendritic segments from cFos+ and neighboring cFos– PL→BLA neurons were targeted for detailed 3D spine morphometric analyses (Figure 4, Supplemental Figures 6, 7, 8). We first investigated the impact of activation by comparing dendritic spines of cFos+ and cFos– neurons in each group, and found little evidence of systematic activity-dependent regulation of dendritic spine density or size in any spine subtype (Figure 4C,D). Next, we investigated the role of stress by comparing the stress group (resilient+susceptible) to the control group. Interestingly, spine density did not differ in any spine subtype for either cFos+ or cFos– neurons, though a stress-induced increase in spine head diameter and length was observed across most spine types and was unexpectedly more pronounced in cFos– neurons (Supplemental Figure 7).

**Fig. 4.**
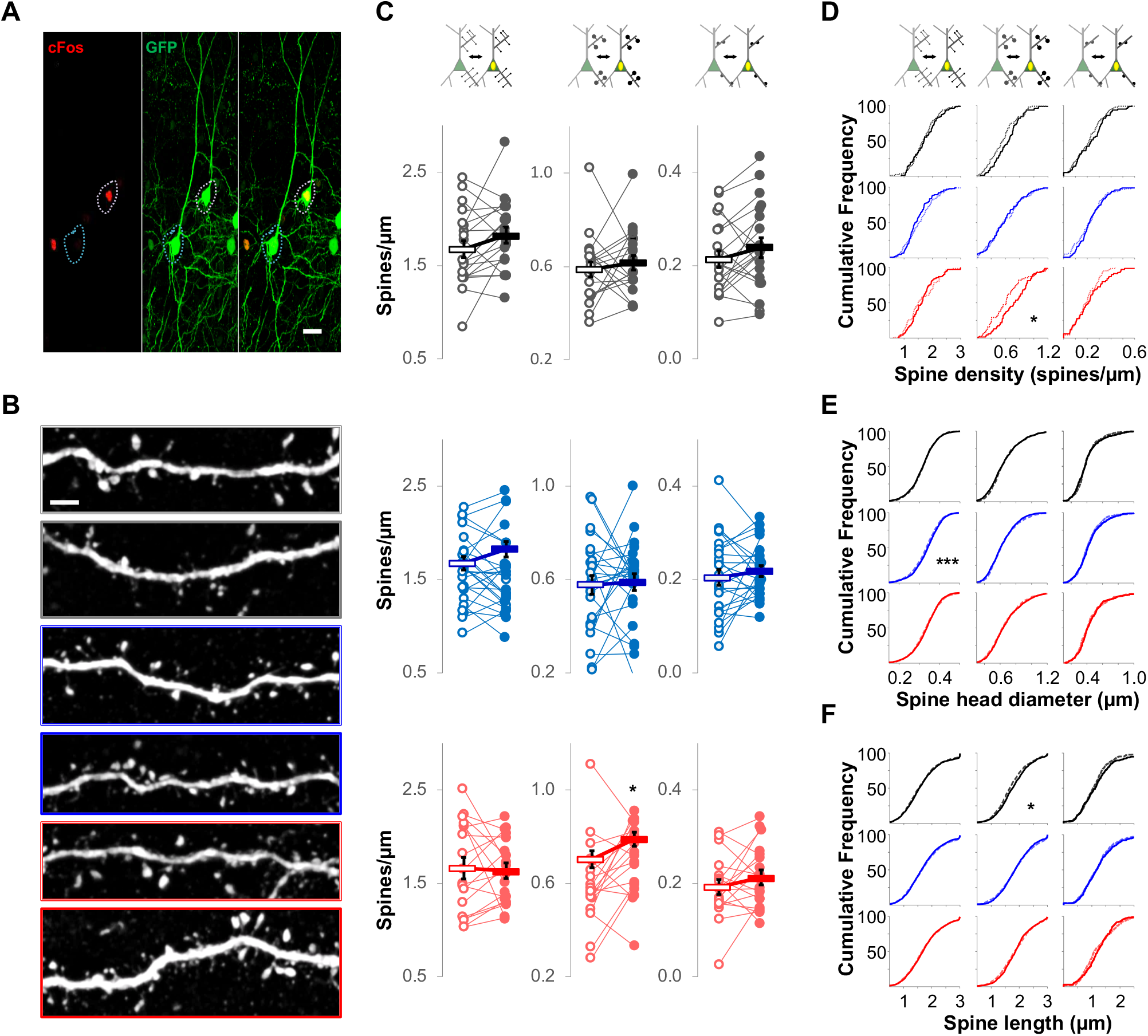
Comparison of dendritic spine density and morphology among cFos+ versus cFos– PL→BLA neurons shows no systematic activity-dependent regulation in any spine subtype. (A) Representative image of neighboring cFos+ and cFos– GFP-expressing neurons (scale bar 20μm). Yellow outline indicates cFos+ neuron. Cyan outline indicates the cFos− neuron. Most cFos+ neurons analyzed had a close neighbor matched to Bregma and layer, enabling paired comparisons. (B) Representative images of apical and basal dendrites from cFos+ neurons (scale bar 3μm; gray outline from control; blue outline from resilient; red outline from susceptible). Single width outlines indicate basal dendrites and double width outlines indicate apical dendrites (C-F) Spine subtypes are separated by column and indicated by the cartoon at the top. cFos+ and cFos– neurons are indicated in the cartoon by yellow oval or no oval, respectively, in the green soma. Spine subtype is shown by length and size of protuberances in cartoon. Thin spines are long and small, mushroom spines are large and long, and stubby spines are short. Black represents data from controls, blue from resilient, and red from susceptible. For all distributions, the reported p-value represents K-S test with Bonferroni correction. (C) First column shows comparison for thin spines, second column for mushroom spines, and third column for stubby spines. Shown are mean densities of paired (neighboring) cFos– (open circles) and cFos+ neurons (closed circles). Bars represent the average of all neurons in the group. Among comparisons, the only significant difference identified was a higher mushroom spine density in cFos+ versus neighboring cFos– neurons from susceptible mice (two-tailed paired T-test, C thin p=0.08, R thin p=0.39, S thin p=0.79, C mushroom p=0.50, R mushroom p=0.81, S mushroom p=0.03, C stubby p=0.18, R stubby p=0.38, S stubby p=0.26). Spine density (D), spine head diameter (E) and spine length (F) were compared for each spine type and group using cumulative distributions frequencies (CDF). Dotted lines represent pooled data from all cFos– neurons in that group, while thick lines represent pooled data from all cFos+ neurons in the group. Dendrites of cFos+ neurons have significantly higher mushroom spine densities than dendrites of cFos– neurons in susceptible mice (p=0.0441). No other significant differences in densities were observed in cFos– versus cFos+ neurons (C thin p=0.544, R thin p=0.171, S thin p=5889, C mushroom p=0.0762, R mushroom p=0.727, C stubby p=0.201, R stubby p=0.256, S stubby p=0.345). For spine head diameter, only thin spines from cFos+ neurons of resilient mice were significantly larger than their counterparts from cFos– neurons (p=0.0008). For spine length, only mushroom spines of cFos+ neurons of control mice were significantly longer than their cFos– counterparts (p=0.028). C=control; R=resilient; S=Susceptible; *p<0.05; ***P<0.001

Despite no stress-associated differences in spine density, susceptible mice had significantly higher mushroom spine density than resilient mice, and this difference was specific to basal dendrites of cFos+ PL→BLA neurons (Figure 5, Supplemental Figure 8). No differences were found in the spine densities of cFos– neurons. Resilient and susceptible mice had larger thin spine head diameters and lengths than controls across cFos+ and cFos– neurons, but the magnitude was higher in susceptible mice in several instances. Additionally, susceptible mice had significantly larger mushroom spine size than resilient mice, though neither group differed significantly from control.

**Fig. 5.**
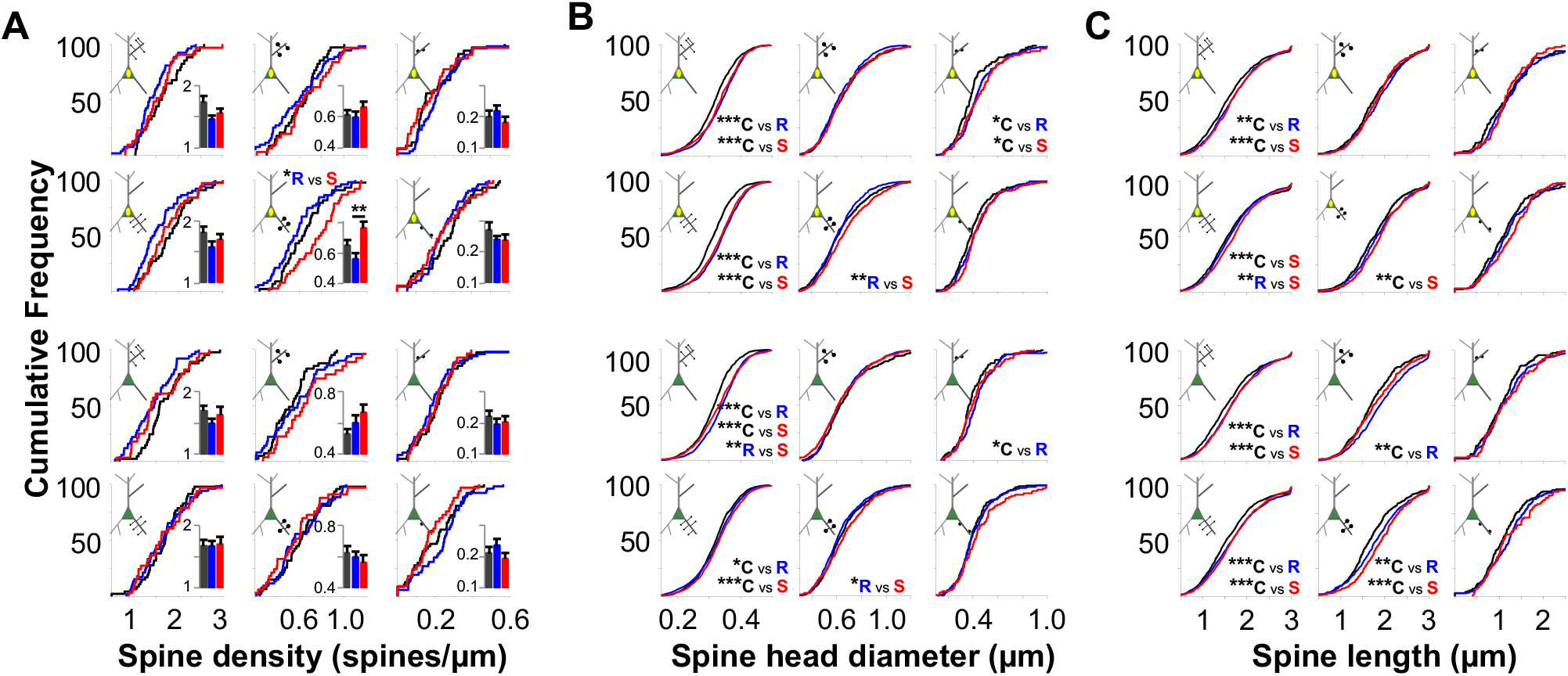
Dendritic spine differences likely represent a combination of preexisting structural differences and differences in stress-induced plasticity, but overall suggest higher excitatory drive in PL→BLA neurons of susceptible mice. (A) Representative image of neighboring cFos+ and cFos– GFP-expressing neurons (scale bar 20μm). Yellow outline indicates cFos+ neuron. Cyan outline indicates the cFos− neuron. Most cFos+ neurons analyzed had a close neighbor matched to Bregma and layer, enabling paired comparisons. (B) Representative images of apical and basal dendrites from cFos+ neurons (scale bar 3μm; gray outline from control; blue outline from resilient; red outline from susceptible). Single width outlines indicate basal dendrites and double width outlines indicate apical dendrites (C-F) Spine subtypes are separated by column and indicated by the cartoon at the top. cFos+ and cFos– neurons are indicated in the cartoon by yellow oval or no oval, respectively, in the green soma. Spine subtype is shown by length and size of protuberances in cartoon. Thin spines are long and small, mushroom spines are large and long, and stubby spines are short. Black represents data from controls, blue from resilient, and red from susceptible. For all distributions, the reported p-value represents K-S test with Bonferroni correction. (C) First column shows comparison for thin spines, second column for mushroom spines, and third column for stubby spines. Shown are mean densities of paired (neighboring) cFos– (open circles) and cFos+ neurons (closed circles). Bars represent the average of all neurons in the group. Among comparisons, the only significant difference identified was a higher mushroom spine density in cFos+ versus neighboring cFos– neurons from susceptible mice (two-tailed paired T-test, C thin p=0.08, R thin p=0.39, S thin p=0.79, C mushroom p=0.50, R mushroom p=0.81, S mushroom p=0.03, C stubby p=0.18, R stubby p=0.38, S stubby p=0.26). Spine density (D), spine head diameter (E) and spine length (F) were compared for each spine type and group using cumulative distributions frequencies (CDF). Dotted lines represent pooled data from all cFos– neurons in that group, while thick lines represent pooled data from all cFos+ neurons in the group. Dendrites of cFos+ neurons have significantly higher mushroom spine densities than dendrites of cFos– neurons in susceptible mice (p=0.0441). No other significant differences in densities were observed in cFos– versus cFos+ neurons (C thin p=0.544, R thin p=0.171, S thin p=5889, C mushroom p=0.0762, R mushroom p=0.727, C stubby p=0.201, R stubby p=0.256, S stubby p=0.345). For spine head diameter, only thin spines from cFos+ neurons of resilient mice were significantly larger than their counterparts from cFos– neurons (p=0.0008). For spine length, only mushroom spines of cFos+ neurons of control mice were significantly longer than their cFos– counterparts (p=0.028). C=control; R=resilient; S=Susceptible; *p<0.05; ***P<0.001

In summary, we identified complex and unique PL→BLA dendritic spine morphological signatures in susceptible versus resilient mice that cannot be explained by activity-dependent regulation (Figure 6). They are also unlikely to be solely due to stress-induced plasticity given that most differences identified specifically in susceptible versus resilient comparisons are related to mushroom spines, which are generally considered to be the most stable subtype[8, 61, 62]. Thus, a higher number and larger size of PL→BLA mushroom spines might represent the structural substrate for aberrant overexcitability of this pathway in susceptible mice.

**Fig. 6.**
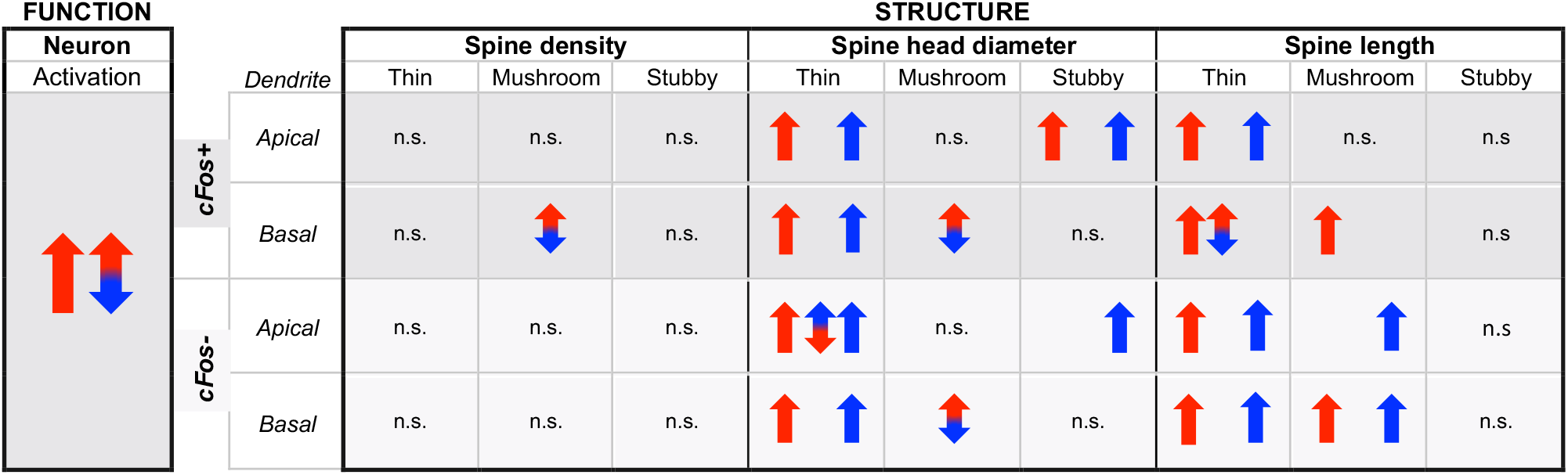
Dendritic spine differences likely represent a combination of preexisting structural differences and differences in stress-induced plasticity, but overall suggest higher excitatory drive in PL→BLA neurons of susceptible mice. (A) Representative image of neighboring cFos+ and cFos– GFP-expressing neurons (scale bar 20μm). Yellow outline indicates cFos+ neuron. Cyan outline indicates the cFos− neuron. Most cFos+ neurons analyzed had a close neighbor matched to Bregma and layer, enabling paired comparisons. (B) Representative images of apical and basal dendrites from cFos+ neurons (scale bar 3μm; gray outline from control; blue outline from resilient; red outline from susceptible). Single width outlines indicate basal dendrites and double width outlines indicate apical dendrites (C-F) Spine subtypes are separated by column and indicated by the cartoon at the top. cFos+ and cFos– neurons are indicated in the cartoon by yellow oval or no oval, respectively, in the green soma. Spine subtype is shown by length and size of protuberances in cartoon. Thin spines are long and small, mushroom spines are large and long, and stubby spines are short. Black represents data from controls, blue from resilient, and red from susceptible. For all distributions, the reported p-value represents K-S test with Bonferroni correction. (C) First column shows comparison for thin spines, second column for mushroom spines, and third column for stubby spines. Shown are mean densities of paired (neighboring) cFos– (open circles) and cFos+ neurons (closed circles). Bars represent the average of all neurons in the group. Among comparisons, the only significant difference identified was a higher mushroom spine density in cFos+ versus neighboring cFos– neurons from susceptible mice (two-tailed paired T-test, C thin p=0.08, R thin p=0.39, S thin p=0.79, C mushroom p=0.50, R mushroom p=0.81, S mushroom p=0.03, C stubby p=0.18, R stubby p=0.38, S stubby p=0.26). Spine density (D), spine head diameter (E) and spine length (F) were compared for each spine type and group using cumulative distributions frequencies (CDF). Dotted lines represent pooled data from all cFos– neurons in that group, while thick lines represent pooled data from all cFos+ neurons in the group. Dendrites of cFos+ neurons have significantly higher mushroom spine densities than dendrites of cFos– neurons in susceptible mice (p=0.0441). No other significant differences in densities were observed in cFos– versus cFos+ neurons (C thin p=0.544, R thin p=0.171, S thin p=5889, C mushroom p=0.0762, R mushroom p=0.727, C stubby p=0.201, R stubby p=0.256, S stubby p=0.345). For spine head diameter, only thin spines from cFos+ neurons of resilient mice were significantly larger than their counterparts from cFos– neurons (p=0.0008). For spine length, only mushroom spines of cFos+ neurons of control mice were significantly longer than their cFos– counterparts (p=0.028). C=control; R=resilient; S=Susceptible; *p<0.05; ***P<0.001

### Chemogenetic inhibition of the PL→BLA pathway during or prior to ASDS shifts the population response toward resilience

To test the hypothesis that PL→BLA overexcitability during ASDS mediates the susceptible behavioral response, we used an intersectional chemogenetic approach to deliver an inhibitory DREADD to this pathway (Figure 7A-C). In our first experiment, CNO was administered only once, 30 min prior to ASDS. CNO treatment in animals with histologically confirmed injections was found to decrease cFos both overall in the PL, as well as specifically in PL→BLA neurons (Figure 7D-F). Consistent with our hypothesis, saline injected mice showed a typical pattern of social avoidance, while CNO injected defeated mice did not differ from controls in their interaction with a novel mouse or time spent in corner zones (Figure 7G). In stereotactically injected mice with no DREADD expression, both saline and CNO groups showed social avoidance.

**Fig. 7.**
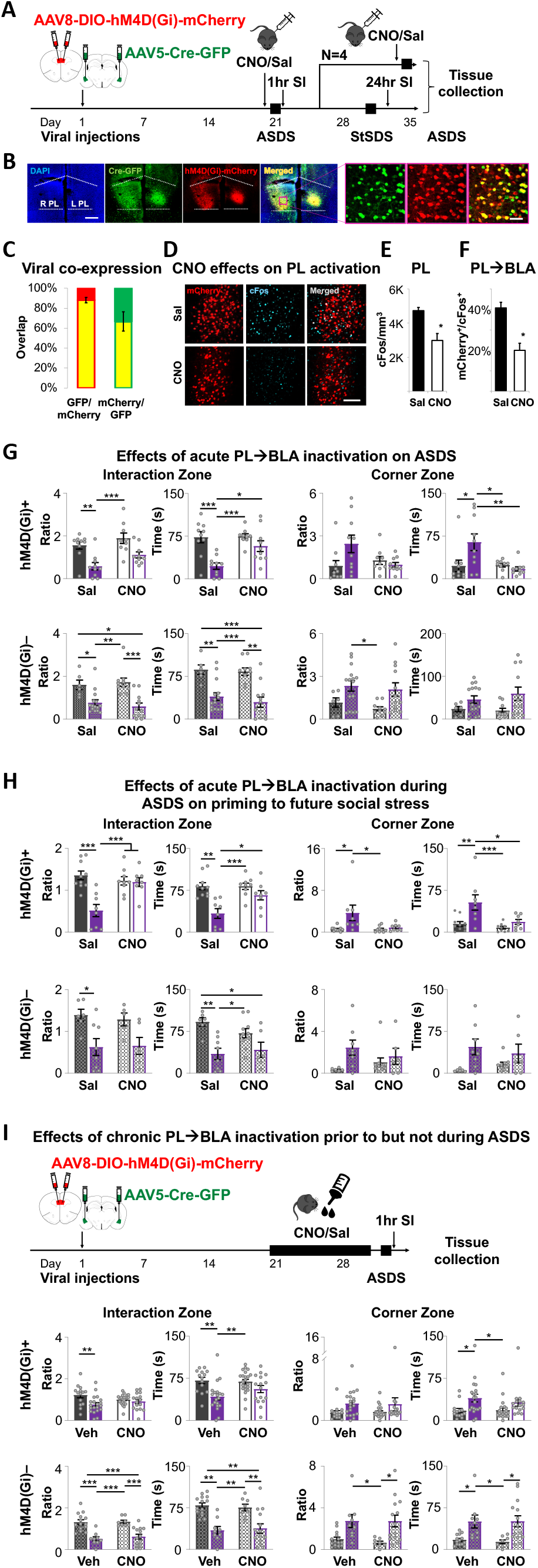
Chemogenetic inhibition of PL→BLA during and prior to ASDS shifts the population response toward resilience. (A) Experimental timeline for acute chemogenetic inhibition. Mice were stereotactically injected with a retrograde Cre-GFP-carrying virus in BLA bilaterally and a Cre-dependent hM4Di-mCherry-carrying virus in PL bilaterally. Following three weeks recovery, mice were injected intraperitoneally (i.p.) with either saline or CNO dissolved in saline 30 minutes prior to ASDS and SI. One week later, a small subset of mice underwent another ASDS and SI in the presence of CNO or saline, in order to collect brains for studies looking at CNO effects on cFos and the fidelity and efficiency of the intersectional chemo-genetic approach. Remaining mice were subjected to StSDS on day 28 and were tested on SI 24 hours later. All mice were then sacrificed and brains collected for histological examination of viral injections. (B) Representative PL image (scale bar 0.5mm) of intersectional viral approach. Inset (pink outline) shows a portion of the PL at higher resolution (scale bar 200μm). Colocalized hM4D(Gi)-mCherry/Cre-GFP appear yellow in merged image on right. (C) Intersectional approach showed high fidelity [88.51%±5.07 of hM4D(Gi) puncta (red) colocalized with Cre puncta (green)] and moderately high efficiency [67.86%±19.63% of Cre puncta colocalized with hM4D(Gi) puncta]. (D) Representative image (scale bar 200 μm) of hM4D(Gi)-mCherry (red) and cFos (cyan) after i.p. saline or CNO 30 min prior to ASDS and SI, followed by immediate sacrifice. (E) Significantly lower cFos density (two-tailed unpaired T-test, p=0.018) was observed in the PL of CNO compared to saline injected mice. (F) Significantly fewer cells expressing hM4D(Gi) (red) co-expressed cFos (cyan) in CNO compared to saline injected mice (two-tailed unpaired T-test, p=0.0015). (G-I) For all group comparisons, two-way ANOVA with post-hoc Tukey correction were used to test drug by stress interactions (DrugXStress), main stress effects (stress), main drug effects (drug), and individual differences between control (C) and defeated (D) mice treated with saline (sal) versus CNO. (G) Effects of PL→BLA inhibition during ASDS. SI ratio, corner ratio, and absolute time in the interaction zone and corner zones was compared between hM4D(Gi)+ mice (defined as histologically verified targeting of all four injection sites) in sal versus CNO groups. In contrast to saline-injected mice, which showed a typical pattern of social avoidance following ASDS, CNO injected mice subjected to ASDS had control-level SI ratios [DrugXStress F(1,35)=0.45, p=0.60, stress F(1,35)=36 p<0.0001, drug F(1,35)=8.6 p=0.026, sal C v sal D p=0.0034, sal D v CNO C p=0.0001], interaction times [DrugXStress F(1,35)=7.1, p=0.040, stress F(1,35)=29 p<0.0001, drug F(1,35)=8.5 p=0.025, sal C v sal D p=0.0003, CNO C v sal D p=0.0003, sal D v CNO D p=0.017], time in corner zone [DrugXStress F(1,35)=13, p=0.015, stress F(1,35)=6.6 p=0.074, drug F(1,35)=11 p=0.022, sal C v sal D p=0.02, CNO C v Sal D p=0.033, sal D v CNO D p= 0.0066), and corner ratios [DrugXStress F(1,35)=12, p=0.026, stress F(1,35)=5.4 p=0.13, drug F(1,35)=4.1 p=0.18], with three of these four indices showing significant DrugXStress interaction. In contrast, for mice with failed intersectional approach [hM4D(Gi)–], both saline- and CNO-injected defeated mice showed the typical social avoidance pattern of decreased SI ratios [DrugXStress F(1,39)=0.96, p=0.45, stress F(1,39)=41 p<0.0001, drug F(1,39)=0.031 p=0.88, sal C v sal D p=0.024, sal C v CNO D p=0.0056, CNO C v sal D p=0.001, CNO C v CNO D p=0.0002] and interaction time [DrugXStress F(1,39)=0.18, p=0.71, stress F(1,39)=48 p<0.0001, drug F(1,39)=0.81 p=0.43, sal C v sal D p=0.0029, sal C v CNO D p=0004, CNO C v sal D p=0.001, CNO C v CNO D p=0.0001], and increased corner ratios [DrugXStress F(1,39)=0.10, p=0.82, stress F(1,39)=19 p=0.0036, drug F(1,39)=1.4 p=0.40, CNO C v sal D p=0.021] and corner times [DrugXStress F(1,39)=1.1, p=0.48, stress F(1,39)=14 p=0.013, drug F(1,39)=0.40 p=0.67], with all comparisons showing a main stress effect and no significant DrugXStress interaction. (H) Effects of PL→BLA inhibition during ASDS on priming to susceptibility on subsequent StSDS. Sal versus CNO refers to administration during ASDS. No drug was administered during StSDS. CNO administration during ASDS protected hM4D(Gi)+ mice from stress-priming effects as evidenced by control-level SI ratios [DrugXStress F(1,31)=20, p=0.0013, stress F(1,31)=23 p=0.0006, drug F(1,31)=9.2 p=0.022, sal C v sal D p<0.0001, CNO C v sal D p=0.0009, sal D v CNO D p=0.002], interaction times [DrugXStress F(1,31)=8.5, p=0.03, stress F(1,31)=33 p<0.0001, drug F(1,31)=8.3 p=0.032, sal C v sal D p=0.002, CNO C v sal D p=0.0003, sal D v CNO D p=0.027], corner times [DrugXStress F(1,31)=7.1, p=0.059, stress F(1,31)=22 p=0.0017, drug F(1,31)=16 p=0.0061, sal C v sal D p=0.0032, CNO C v sal D p=0.0006, sal D v CNO D p=0.0134], and corner ratios [DrugXStress F(1,31)=8.9, p=0.056, stress F(1,31)=14 p=0.02, drug F(1,31)=9.2 p=0.052, sal C v sal D p=0.02, CNO C v sal D p=0.023], with significant DrugXStress interactions in two of the four indices and trends toward significance in the other two indices. In contrast,no DrugXStress interactions were seen for hM4D(Gi)– mice in SI ratios [DrugXStress F(1,21)=0.36, p=0.72, stress F(1,21)=39 p=0.0014, drug F(1,21)=0.11 p=0.84, sal C v sal D p=0.038], interaction times [DrugXStress F(1,21)=4, p=0.19, stress F(1,21)=40 p=0.0002, drug F(1,21)=0.89 p=0.53, sal C v sal D p=0.0036, sal C v CNO D p=0.023, CNO C v CNO D p=0.048], corner ratios [DrugXStress F(1,21)=5.1, p=0.20, stress F(1,21)=16 p=0.029, drug F(1,21)=0.038 p=0.91], or corner time [DrugXStress F(1,21)=3.2, p=0.30, stress F(1,21)=23 p=0.008, drug F(1,21)=0.0064 p=0.96], but significant main stress effects were observed in all four indices, indicating ASDS-priming effects in both saline and CNO groups. Experimental timeline for chronic chemogenetic inhibition of PL→BLA. Same intersectional chemogenetic approach and post-hoc histological inspection of injection sites was performed as described above. Following three week recovery, mice were given either CNO or vehicle-treated (aspartame) drinking water *ad libitum* for 10 days. On Day 11, regular drinking water was provided in the morning and mice underwent ASDS and SI in the afternoon. As in the case of acute CNO administration, vehicle-treated hM4D(Gi)+ mice showed the typical pattern of social avoidance following ASDS, but CNO-treated mice showed partial protection from the phenotypic decreased SI ratios [DrugXStress F(1,68)=4.7, p=0.056, stress F(1,68)=11 p=0.0044, drug F(1,68)=0.18 p=0.70, sal C v sal D p=0.0069] and interaction times [DrugXStress F(1,68)=2.3, p=0.16, stress F(1,68)=17 p=0.0003, drug F(1,68)=1.2 p=0.30, sal C v sal D p=0.0032, sal D v CNO C p=0.0019], and increased corner zone time [DrugXStress F(1,68)=0.71, p=0.45, stress F(1,68)=13 p=0.0021, drug F(1,68)=0.44 p=0.55, sal C v sal D p=0.041, sal D v CNO C p=0.023], and corner ratio [DrugXStress F(1,68)=0.060, p=0.83, stress F(1,68)=7.4 p=0.022, drug F(1,68)=0.0001 p=0.99]. None of these indices reached significance in DrugxStress interactions, but a trend was observed in two of them. In contrast, hM4D(Gi)– mice showed a typical pattern of ASDS-induced social avoidance irrespective of CNO treatment, with main stress effects in both vehicle and CNO-treated mice and no DrugxStress interactions on SI ratio [DrugXStress F(1,41)=0.20, p=0.68, stress F(1,41)=50 p<0.0001, drug F(1,41)=0.32 p=0.60, sal C v sal D p<0.0001, sal C v CNO D p<0.0001, CNO C v sal D p=0.0003, CNO C v CNO D p=0.0007], interaction time [DrugXStress F(1,41)=0.33, p=0.62, stress F(1,41)=43 p<0.0001, drug F(1,41)=0.001 p=0.98, sal C v sal D p=0.0003, sal C v CNO D p=0.0002, CNO C v sal D p=0.0048, CNO C v CNO D p=0.0045], corner time [DrugXStress F(1,41)=0.094, p=0.81, stress F(1,41)=30 p=0.0001, drug F(1,41)=0.060 p=0.85, sal C v sal D p=0.037, sal C v CNO D p=0.012, CNO C v sal D p=0.045, CNO C v CNO D p=0.02], and corner ratios [DrugXStress F(1,41)=0.23 p=0.72, stress F(1,41)=29 p=0.0002, drug F(1,41)=0.24 p=0.71, sal C v CNO D p=0.027, CNO C v sal D p=0.045, CNO C v CNO D p=0.022].

To test if the effects of inhibiting the PL→BLA extended to blocking the priming effect of ASDS on future susceptibility, mice were also tested on StSDS in the absence of CNO following a one week recovery (Figure 7H). As expected, mice that had been saline-treated during ASDS showed a priming effect, displaying social avoidance following StSDS. In contrast, the priming effect was completely blocked in mice that had been CNO-treated during ASDS, implicating the PL→BLA in ASDS fear memory acquisition, rather than solely in fear expression.

In our second experiment, we asked if long-term inhibition of the PL→BLA would remain effective in shifting the population response toward resilience. CNO or vehicle was added to the drinking water and provided *ad libitum* for 10 days (Figure 7I, Supplemental Figure 9). On day 11, mice underwent ASDS without CNO administration. Similar to acute CNO administration, chronically CNO-treated defeated mice showed control level social interaction while vehicle treated mice and mice with no DREADD expression showed the typical pattern of social avoidance.

## Discussion

We present data on a novel acute stressor that uniquely, rapidly, and reproducibly classifies male mice into susceptible versus resilient. ASDS mirrors CSDS population distribution of social avoidance, does not induce CSDS-phenotypic long-term maladaptive behaviors, but does prime for increased susceptibility to a future social stressor. This suggests some form of fear memory acquisition occurs despite no changes in baseline behavior. Although the comparison between ASDS and CSDS classification into susceptible versus resilient is limited by this priming effect, the two models likely tap into overlapping behavioral endophenotypes. This is supported by (i.) the striking consistency in distributions of susceptibility and resilience on ASDS versus CSDS; (ii.) all mice that remain resilient after ASDS followed by CSDS were originally classified as resilient on ASDS, and (iii.) resilience on ASDS confers some protection from subsequent CSDS-induced social avoidance.

This novel paradigm was developed to offer a window into preexisting neurocircuit structure/function by enabling histological and morphological investigation of the stress-induced neuronal ensemble prior to major stress-induced circuit reorganization. While synaptic connectivity can be altered on the order of minutes to hours in response to a stimulus[44, 47, 46, 61, 62],, major stress-induced rewiring is unlikely on this timescale. The PL→BLA pathway is a cornerstone of the stress-response and highly implicated in fear memory acquisition[1, 10, 17, 45, 67, 85, 92]. Therefore, the higher recruitment of PL→BLA neurons in susceptible mice during a first-time stressor likely occurs as a function of pre-existing differences in local and/or long-range neural inputs. This is supported by the overall pattern of higher density and size of PL→BLA mushroom spines in susceptible mice, the spine subtype known for both stability and highest density of excitatory receptors[8, 43, 42, 61, 62, 96, 97].

Importantly, differences in mushroom spines were restricted to susceptible versus resilient comparisons, with no differences observed between control versus resilient or control versus susceptible comparisons, or between control and the combined (resilient+susceptible) stress group. In contrast, thin spine head diameter and length showed an indiscriminate stress-induced increase in both resilient and susceptible mice across cFos+ and cFos– neurons. This serves as an inherent positive control, since thin spines are well-documented to be more plastic than mushroom spines[5, 7, 8, 9, 11, 12, 24, 28, 33, 36, 43, 42, 47, 61, 62, 86, 94] and show rapid changes in response to systemic stress hormones[38, 48, 62].

Using slice physiology and a different stress model, Wang et al. showed rapid and opposing changes in excitatory transmission in cFos+ PL neurons from susceptible versus resilient mice[93]. Spine turnover and density changes have also been observed on this timescale[77], though these seem to be restricted to thin spines[62]. Furthermore, *in vivo* observations of dendritic spine formation indicate that large spines form by enlargement of preexisting smaller spines[8, 59]. However, rapid stress-induced maturation of thin spines into mushroom spines in susceptible mice is unlikely in the setting of no compensatory decrease in thin or stubby spine density. Interestingly, PL neuron stress-induced effects have mostly been observed in apical dendrites[9, 31, 37, 65], while our mushroom spine density difference was exclusive to basal dendrites. Our experimental design does not allow precise delineation of the relative contribution of preexisting structural differences versus stress-induced plasticity, but, collectively, observed mushroom spine differences most likely represent preexisting connectivity differences, while observed thin spine changes most likely represent stress-induced plasticity. Confirming this will require methods that can monitor these structures longitudinally.

The source of input to the more numerous and larger mushroom spines on PL→BLA neurons from susceptible mice is unknown at this time. There is some evidence the amygdala may preferentially synapse onto basal dendrites of PL neurons[56] and activation of the reciprocal BLA→PL circuit reduces social interaction[26]. Thus, an intriguing possibility for observed structure/function differences could be an aberrant positive feedback loop between the PL and BLA.

Irrespective of the origin of these inputs, our findings implicate PL→BLA overexcitability during a first-time stressor as a biomarker of trait vulnerability. In accordance with this hypothesis, and consistent with the literature implicating this circuit in fear expression[45, 71, 87], we found inhibiting this pathway during ASDS blocked stress-induced social avoidance selectively in DREADD expressing mice, with stress by drug interactions significant in three of four behavioral indices measured (interaction time, corner ratio, and corner time). Additionally, inactivation of PL→BLA during ASDS also blocked ASDS-induced priming to future stress, which is consistent with the literature implicating this pathway also in fear memory consolidation[2, 18, 25, 29, 74, 87, 90].

Given that our goal here was to investigate potential targets for prevention rather than treatment, it is important to ascertain if long-term manipulations of a circuit lead to habituation and/or compensatory circuit mechanisms. We therefore also tested the effects of chronic PL→BLA inhibition by CNO administration in drinking water during the 10 days prior to ASDS. Our results show no evidence of circuit habituation or compensatory mechanisms. Chronic CNO treatment was equally effective in blocking ASDS-induced social avoidance, and this could not be explained by previously reported off-target CNO-drug effects[32, 41, 57, 58, 70, 75] given that DREADD non-expressing mice showed expected pattern of social avoidance. Important next steps include ascertaining if such chronic manipulations can “rewire” the PL→BLA circuit to induce a lasting resilient state, identifying the structural changes during such rewiring, assessing the duration of preventative protection against stress, and extending findings to female mice.

## ACKNOWLEDGEMENTS

This research was supported by a 2015 NARSAD Young Investigator Award (DD), R01MH111918 (DD) and funding from the Departments of Pediatrics, Environmental Medicine & Public Health and Neuroscience at the Icahn School of Medicine at Mount Sinai. We thank Drs. Eric Nestler, Scott Russo, Patrizia Casaccia and Hiro Morishita helpful suggestions on experimental design. We also thank Kristin Anderson for her assistance in formatting this manuscript for online publication.

## AUTHOR CONTRIBUTIONS

YSG, CF and DD designed experiments, YSG, CF, AM, RW, TZ and DD performed experiments, YSG, CF, TZ and DD analyzed the data. YSG, AM and DD wrote the manuscript. All authors read and approved the manuscript.

## Extended Methods

### Animals

For experimental mice, 7-12 week old C57BL/6J male mice were purchased from Jackson Laboratory and group housed (4 per cage) in standard mouse cages on a 12 hour light cycle (light on 07:00-19:00) with ad libitum food and water. For aggressors, sexually experienced retired breeder CD1 male mice were purchased from Charles River Laboratories and housed individually with *ad libitum* food and water in standard cages. After allowing three to seven days acclimation, CD1 mice were screened for aggression over the course of three days by introducing an intruder C57BL/6J male mouse into their home cage for five minutes on the first day and three minutes on subsequent days. Only CD1 mice that 1) expressed aggression on each of the three days of screening, 2) exhibited latency to aggression <30 seconds on the second and third day and 3) demonstrated multiple bouts of aggression during the screening session on the third day were used in the social defeat experiments. Non-aggressive CD1 mice were used in the social interaction test (SI). All behavioral experiments were performed between 13:00 and 18:00, i.e. during light on.

All experiments were conducted in compliance with the National Institutes of Health Guidelines for the Care and Use of Experimental Animals approved by the Institutional Animal Care and Use Committee at the Icahn School of Medicine at Mount Sinai.

### Chronic social defeat stress (CSDS)

CSDS was performed as previously described[6, 30, 50, 52, 91]. In brief, experimental C57BL/6J mice were placed into the cage (larger hamster cages was used for CSDS) of a novel aggressive CD1 mouse for 5 min daily for 10 consecutive days. For the remainder of each 24 hour period, a clear perforated barrier was placed between the C57BL/6J mouse and resident aggressor to allow sensory, but not physical, interaction. Control C57BL/6J mice also rotated to a novel mouse’s cage each day for 10 days, but the resident was a non-aggressive conspecific mice. At the end of 10 days, C57BL/6J mice were group-housed with prior cage mates. Each group-housed cage of experimental mice contained either four defeated or four control mice to avoid stress phenotype transfer. On the 11th day, experimental mice were tested in a social interaction test as described below.

### Subthreshold defeat stress (StSDS)

StSDS has been previously described as a submaximal stressor that does not lead to social avoidance[30, 50, 51]. This test is therefore used to assess increased susceptibility in response to various manipulations. Experimental C57BL/6J mice were placed into the cages of three aggressive CD1 mice for 5 min each, with 15 min rest in a clean cage between aggressive sessions. Control mice were placed into the cages of three novel non-aggressive conspecific mice. Experimental and control mice were then returned to their home cages with their prior cage mates and tested on SI 24 hours later.

### Social interaction test (SI)

SI testing was performed identically for CSDS, StSDS and ASDS. C57BL/6J mice were placed into an opaque Plexiglas open-field arena (42×42×42cm^3^) with a removable wire-mesh enclosure secured in clear Plexiglas (10 cm (w) × 6.5 cm (d) × 42 cm (h)) placed against the middle of one of the inner walls of the arena. Mice were allowed to explore the arena for 150 sec. The mouse was then briefly (30 sec) removed from the arena and a novel non-aggressive CD1 “target” mouse was placed in the wire-mesh enclosure. The C57BL/6J mouse was then returned to the arena and allowed to explore for another 150 sec. Open-field arenas and wire-mesh enclosures were wiped with a cleaning solution containing 70% ethyl alcohol between tests. All tests were conducted during the lights on period (between 13:00 and 19:00) under red light conditions. A video-tracking system (Ethovision 3.0, Noldus Information Technology) was used to record the movements of the C57BL/6J mouse during the “target absent” and “target present” periods. The total time spent in the “interaction zone” (8 cm wide corridor surrounding the wire-mesh enclosure) and “corner zones” (9 cm × 9 cm in the two corners on the opposite wall from the wire-mesh), as well as the total distance traveled and average velocity were recorded during both periods. SI and corner ratios were calculated as time spent in the respective zone when target is present divided by time spent in the zone when target is absent.

For experiments establishing the ASDS model, “resilience” was defined as SI ≥ 1 and “susceptibility” was defined as SI ratio<1 in accordance with the conventional classification for CSDS[50]. For experiments looking at PL*to*BLA activation and dendritic spine structure, a multidimensional classifier for resilience and susceptibility was used. These criteria were designed for three reasons. First, they exclude the weakly avoidant or socially “indifferent” behavior observed in a subset of both control and defeated mice. Secondly, using this classification system on ASDS-defeated mice has high sensitivity and specificity for mice remain resilient in a subsequent CSDS (see RESULTS), indicating that using a multidimensional classifier taps into a homogenous trait-resilience phenotype. Third, because we hypothesized putative pre-existing functional and structural differences to be very small given that they represent individual variability within a naïve population. These criteria were used to enhance the homogeneity of behavioral groups. All animals with an average velocity less than 1.8 cm/s were excluded. This population reflects 7.5% (N=27 of 359) of defeated mice with freezing rather than avoidant behavior. This behavior is exceedingly rare in control mice (1.3%, N=2 of 149). Because these mice can freeze anywhere within the arena when a target mouse is present, they occasionally freeze at the edge of the interaction zone leading to an artificially elevated SI ratio. In the PL*to*BLA activation and dendritic spine structure study, this population reflected 2.2% of defeated mice (N=1 of 46). Resilience was then defined as meeting all three of the following criteria: SI ≥ 1, absolute time interacting with novel target mouse ≥60 sec, and absolute time spent in corners when novel target mouse present < 30 sec. Susceptibility was defined as requiring both of the following criteria: SI ratio less than 0.6 and absolute time spent interacting with target mouse less than 50 sec. In our combined 17 experiments establishing the ASDS protocol, 68% (N=101 of 149) of control mice met the resilience criteria and only 5.4% (N=8 of 149) met the susceptibly criteria. In contrast, if a simple SI ratio below 1 is used to classification, 24% of control mice (N=35 of 149) would meet susceptibility criteria. Similarly, in the PL*to*BLA activation and structure study, 88% (N=13 of 16) of controls met resilience criteria and only 7% (N=1 of 16) met susceptibility criteria. These criteria are therefore useful for capturing the effects of social defeat without the confounder of weakly avoidant or socially indifferent mice.

Using this multidimensional classifier, resilience represents 26% (N=92 of 359) of the defeated population in our combined 17 experiments establishing the ASDS protocol, and 17% (N=8 of 46) of the defeated population in the three experiments examining PL*to*BLA activation and morphological differences. Susceptible animals represent 32% (N=115 of 359) and 46% (N= 21 of 46) in these populations respectively. Because the PL*to*BLA activation and structure difference study also required animals to have adequate stereotactic injections targeting the BLA and there were only eight resilient mice, one exception was made: one mouse meeting only two of the three criteria (absolute time interacting with novel target mouse and corner time) for resilience and had proper targeting of the BLA was included in the resilient group. In total, seven mice per group (control, resilient, susceptible) fit these criteria, had adequate targeting of the BLA, and were therefore included in post-hoc analyses.

### Sucrose preference

During sucrose preference testing, mice were individually housed to enable quantification of consumed fluids. The standard water bottles were removed and replaced with two 50 mL conicals with sipper stops, one containing plain water and the other containing 1% sucrose in water. Food continued to be provided *ad libitum*. At the end of a 24 hour period the amount of fluid consumed from each bottle was manually recorded and mice were returned to their home cages with their previous cage mates.

### Open-field test

Mice were placed in an empty opaque Plexiglas open-field arena (42×42×42cm^3^) allowed to explore for 5 minutes. Exploratory behavior was recorded with video-tracking software (Ethovision 3.0, Noldus Information Technology). Time spent in the center area (10×10cm) and total distance traveled were calculated.

### Elevated plus maze

Mice were placed in the center of a plus maze apparatus made from black Plexiglas with two open arms and two closed arms [12cm (w) × 50cm (d) × 0.5cm (h)] placed 1 m above floor level. Mice were allowed to explore freely for five minutes and their movements were recorded using video-tracking software (Ethovision 3.0, Noldus Information Technology). Time spent in the open arms, closed arms and total distance traveled were calculated.

### Stereotactic viral injections

#### PL→BLA activation and dendritic spine structure

AAV5-hSyn-eGFP (cat#AV-5-1696, University of Pennsylvania Vector Core) was injected into the BLA to label PL→BLA neurons. Mice were anesthetized with a cocktail of ketamine (100 mg/kg) and xylazine (10 mg/kg) and placed in a stereotactic frame using non-penetrating ear bars (Harvard Apparatus). The scalp was sanitized with betadine and the skull was the exposed via a single midline incision using a sterile scalpel. A small burr whole was made in the scalp overlying the BLA using an electric dental drill. A Hamilton syringe (cat#7641-01, Hamilton Company) fitted with a 26 gauge needle (cat#7758-02, Hamilton Company) was slowly lowered unilaterally into the BLA (from Bregma: medio-lateral +/−3.4, anterio-posterior −1.1, dorso-ventral −5.0, angle 0°. A total of 0.6 *μ*l of AAV5-hSyn-eGFP was injected at a rate of 0.1 *μ*l/min and the syringe was left in place for 10 min to minimize the backdraft of virus. The syringe was then slowly removed at a rate of 0.5 mm/min. Mice were allowed to recover and fully express the virally-delivered eGFP for a minimum of 18 days.

Viral injections were performed unilaterally in a counterbalanced fashion, with half the mice injected in the left BLA and half the mice in the right BLA. All mice that underwent viral injections and behavioral characterization were checked for adequate BLA targeting, which we defined as dense labeling in both the BLA in the injected hemisphere as well as in the contralateral BLA. A total of 48 mice were injected in 3 sets of 16 mice per group. 21 of the injected mice (n= 7 control, 7 resilient, 7 susceptible) met both the viral localization criteria and as well as the behavioral classification described in the “Social Interaction test (SI)” section above, and were therefore included in the activation and dendritic spine study.

#### Chemogenetic inhibition of PL→BLA pathway

An intersectional chemogenetic approach was employed to specifically target the inhibitory Designer Receptor Exclusively Activated by Designer Drug (DREADD) hM4Di to BLA-projecting PL neurons. Animals were anesthetized and prepped as described above. The 0.6 *μ*l of AAV5-hSyn-Cre-GFP (cat#AV-5-PV1848, University of Pennsylvania Vector Core) was injected into the BLA bilaterally, as described above. 0.3 *μ*l AAV8-hSyn-DIO-hM4Di-mCherry (cat#44362, Addgene) were bilaterally infused into the PL (from Bregma: medio-lateral +/−0.8, anterio-posterior +2.3, dorso-ventral −2.3, angle 11°). All animals were allowed a minimum of three weeks of recovery and stable viral-mediated gene expression before commencing behavioral experiments. For acute PL→BLA inhibition, a total of 112 mice were injected in four sets of 6-47 mice. For chronic PL→BLA inhibition, a total of 117 mice were injected in three sets of 38-40 mice.

### Drug administration

Acute CNO administration: During ASDS, control and defeat mice were randomly assigned to injection with either clozapine-N-oxide (CNO, 2 mg/kg, dissolved in saline) or saline 30 min prior to ASDS. A subset of defeated mice (n= 4) were then used for assessing the effect of CNO on inhibiting PL→BLA as evidenced by decreased cFos induction. These mice were subjected to the ASDS paradigm again, one week after the initial experiment, followed by immediate sacrifice. These mice received the same injection (CNO or saline) as in the original experiment. Immunohistochemistry for cFos and quantification of percent activation was performed as described below. These mice were also used in immunohistochemical assessment of eficiencty and fidelity of the intersectional approach by quantifying percent GFP-neurons co-expressing mCherry (67.86%±19.63% Mean± SEM) and percent mCherry-neurons co-expressing GFP (88.51%±5.07% Mean± SEM) within the PL, respectively.

The remaining animals (n=108) were allowed one week of recovery, followed by StSDS which was performed in the absence of any drug delivery to test the effects of inhibiting PL→BLA pathway on the observed priming by ASDS to subsequent increased susceptibility to a subthreshold social defeat paradigm.

Following the completion of behavioral experiments, mice were sacrificed and injection localization was evaluated throughout the anterio-posterior axis for both Cre-recombinase expression, as evidence by GFP fluorescence, as well as DREADD expression, as evidenced by mCherry expression. 35 animals (32.4%, control saline n=10, control CNO n=9, defeat saline n=8, defeat CNO n=8) were found to have adequate localization of all four injection sites as evidenced by both localization of GFP signal centered in the BLA bilaterally, as well as mCherry signal primarily or exclusively within the PL bilaterally. 43 animals (38.9%, control saline n=6, control CNO n=10, defeat saline n=15, defeat CNO n=12) did not show any mCherry expression and thus were deemed to have failed the intersectional chemogenetic approach. These animals were used as DREADD negative virally-injected groups to evaluated the effects of CNO on our behavioral paradigm. Finally, 40 mice had either unilateral mCherry expression or expression primarily located in surrounding regions, such as anterior cingulate cortex or infralimbic cortex, and were excluded from analysis.

Chronic CNO administration: For chronic administration, cages housing 4-5 mice were randomly assigned to CNO or vehicle for drinking water. Vehicle was water with 0.005% aspartame dissolved. For the CNO group, CNO was dissolved in vehicle at a concentration of 0.25mg/mL. For both groups 300mL of drinking water was prepared for each cage every three days. Mice were given *ad libitum* access to vehicle or drinking and food for 10 days. Weights of all mice were taken before starting the experiment and immediately before ASDS. Amount of drinking water consumed was calculated daily. On day 9, all mice were video recorded during open field testing. Distance traveled and time spent in the center zone were quantified. On day 11, all mice were tested on ASDS and SI.

Following the completion of behavioral experiments, mice were sacrificed and injection localization was evaluated throughout the anterio-posterior axis for both Cre-recombinase expression, as evidence by GFP fluorescence, as well as DREADD expression, as evidenced by mCherry expression. 72 mice (62.07%, control saline n=14, control CNO n=20, defeat saline n=22, defeat CNO n=16) were found to have adequate localization of all four injection sites as evidenced by both localization of GFP signal centered in the BLA bilaterally, as well as mCherry signal primarily or exclusively within the PL bilaterally. The remaining 44 mice (control saline n=6, control CNO n=10, defeat saline n=15, defeat CNO n=12) were used to explore the off-target effects of CNO administration.

### Tissue collection

For studies evaluating activation of PL→BLA neurons, animals were sacrificed immediately after the social interaction test (i.e. 65 minutes after the initiation of ASDS). Mice were anesthetized with chloral hydrate (1.5 g/kg) and perfused transcardially with 1% paraformaldehyde in 1 M phosphate buffer (PB, pH 7.4) for 1 min followed by 4% paraformaldehyde in 1 M PB (pH 7.4) for 12 min at a rate of 9 mL/min using a peristaltic pump. Brains were post-fixed in their native skulls for 6 hours in the same 4% paraformadehyde at 4° C. Brains were then deskulled and maintained in 0.1% sodium azide in 1 M phosphate buffered saline (PBS) at 4° C.

### Immunohistochemistry

Brains were sectioned at 150 *μ*m on a vibratome (cat#VT1000 S, Leica) to preserve as much of the dendritic arbor as possible, thus enabling the visual tracing of dendritic segments back to their somas. Our immunohistochemistry protocol was optimized for antibody penetration in these thick sections. The entire protocol was performed at room temperature. Tissue was first washed in 0.1M PBS three times for five minutes each. Tissue was then incubated in a blocking solution containing 5% goat serum, 2% bovine serum albumin, 0.2% cold-water fish skin gelatin, 0.6% Triton in PBS for one hour. Sections were incubated in rabbit anti-cFos primary antibody (cat#sc-52, Santa Cruz) diluted 1:500 in 500 μL blocking solution for 24 hours. Following five 10 minute washes in 0.3% Triton PBS, tissue was incubated in Alexa Fluor-conjugated goat anti-rabbit secondary antibody (Alexa Fluor-568, cat#A-11011, or Alexa Fluor-647, cat#A-21244, Life Technology) diluted at 1:400 in 400 μL blocking solution for two hours. Following six 20 minute washes in 0.3% Triton PBS, tissue was re-incubated in the above blocking solution for one hour, then incubated in rabbit anti-GFP primary antibody (cat#A-11122, Life Technology) diluted at 1:500 in 500 μL of blocking solution. Sections were washed as after the first primary antibody, and incubated in Alexa Fluor-488-conjugated secondary antibody (cat#A-11008, Life Technology) diluted at 1:400 in 400 μL of blocking solution. After five additional five minute washes, tissue was incubated in 4’,6-diamidino-2-phenylindole (DAPI) for 5 min, mounted on glass slides in Vecatashield mounting media (cat#H-1000-10, Vector Laboratories), and covered with number 1.5 coverslips.

### Confocal imaging

A Zeiss LSM 510 confocal microscope was used for all experiments. For the PL→BLA activation study, sequential dual-channel 3-dimensional tile scans were acquired of the entire medial prefrontal cortex in each of two brain slices per animal with confirmed BLA targeting of viral injection using a 10X air objective (numerical aperture [NA] 0.3) and the following settings: 8 bit images, averaging of 2, pixel dwell time 1.28 μs, voxel size 1.16×1.16×3.0 *μ*m^3^, 488 nm excitation using 15% power and bandpass acquisition of wavelength 505-550 followed by 561 nm excitation using 25% power and bandpass acquisition of wavelength 566-747. Gain for all images at 520 and 530, respectively.

For dendritic spine imaging, three PL cFos immunoreactive (cFos+) and three PL cFos non-immunoreactive (cFos−) neurons per hemisphere were chosen based on the images acquired at 10X. For each neuron, we aimed to acquire 3-dimensional stacks of two apical and two basal dendrites at 100-150 *μ*m distance from the soma. Z-stacks were acquired using a 100X/1.4NA oil objective with the following settings: 8 bit images, averaging of 4, pixel dwell time 1.61μs, voxel size 0.1×0.1×0.1 *μ*m^3^, 488 nm excitation using 15% power and bandpass acquisition of 505-550.

For chemogenetic experiments, quadruple (or sometimes triple, without DAPI) channel images were acquired to image DAPI (as a nuclear marker for histological identification and mapping), GFP, mCherry, and cFos. DAPI was excited at 405 nm and imaged with a bandpass of 420-480 nm. GFP was excited at 488 nm and acquired using a bandpass of 505-550 nm. mCherry was excited at 561 nm and acquired using a bandpass of 575-615 nm. cFos, here stained using an Alexa Fluor-647 secondary antibody, was excited at 633 nm and imaged using a longpass of 650 nm. For confirmation of injection sites, 4-6 150 *μ*m slices containing the full span of the PL and 4-6 150 *μ*m slices containing the full span of the BLA were mounted sequentially and a low resolution image was acquired using the tiling function and a 5x/0.3NA objective. This allowed rapid inspection of the whole anterio-posterior axis for easy localization of the four injection sites. For CNO effects on cFos expression and fidelity and xx of the intersectional chemogenetic approach, higher resolution images were acquired of the PL using a 25x/0.8NA objective and the following settings: 8 bit images, averaging of 2, pixel dwell time 2.56μs, voxel size 0.1×0.1×0.1 *μ*m^3^.

### Quantification of co-localization patterns

Images were first imported into Fiji[79] for preprocessing. Preprocessing settings were determined by comparing automated puncta identification to manual identification. Settings were optimized until automated identification matched experimenter identification for at least 95% of puncta in three different images. In Fiji, images were segmented into their composite channels (488 and 561 for cFos/GFP colocalization, 633 and 561 for mCherry/cFos, and 488 and 561 for mCherry/GFP), then smoothed via mean filtering, prior to top-hat filtering and object identification using Foci Picker 3D (11). For GFP, images were smoothed, then top-hat filtered (7 × 7 × 4) by pixel before puncta were isolated with Foci Picker 3D. For cFos, images were smoothed, then top-hat filtered (4 × 4 × 3) by pixel prior to puncta isolation with Foci Picker 3D. For mCherry, images were smoothed, then top-hat filtered (5 × 5 × 4) by pixel then puncta were identified using Foci Picker 3D. After preprocessing, all images were saved as three-dimensional 8-bit gray-scale tiffs and exported to MatLab.

Custom code was developed for analyzing three-dimensional colocalization of puncta in MatLab. First, an index was taken quantifying the properties of the puncta, including location, size, and fluorescence intensity. Every puncta from each channel was then compared to puncta from other channels and degree of 3D overlap was recorded. Only puncta with >50% overlap were considered colocalized. The code tallied colocalized and non-colocalized puncta per image.

### Dendritic spine quantification

Dendritic spines were quantified using semi-automated quantification previously described in detail[23], but with some important differences. First, deconvolution was not used here as we found that the background created by GFP-expressing axonal processes within the PL interfered with this image-processing step. Secondly, while the semi-automated program NeuronStudio[78] was used, the higher background in this tissue as compared with single cell microinjections (which can be adequately spaced by the researcher to create non-overlapping cells) required significantly more manual adjustments. However, the experimenter was blind to the experimental group and therefore any bias created by this more manual approach was equal for all three groups. Briefly, each z-stack containing a dendrite was imported into NeuroStudio and the dendrite was automatically detected by the program. Automatic detection of spines in 3-dimentions was then complemented by manual inspection of the dendrite by scrolling up and down within the stack and either deleting spines that did not appear connected to dendrite or inserting spines that were missed by the automatic detection. The size and location of each spine and the length of the respective dendrite was recorded by the program and imported into Excel or MatLab for further analysis. Spine density was calculated by dividing the total number of spines on a segment by the length of their parent dendrite. All dendrites were then averaged by cell. Because of large variability in the number of neurons obtained from each animal, individual cell averages were used to detect differences between groups. Since we collected roughly equivalent numbers of dendrites from control, susceptible, and resilient mice, to maintain statistical rigor, comparison of dendritic morphology between control and defeated mice was performed using all of the dendrites from control mice and a semi-random subset of dendrites from defeated mice. The “defeat” subset was evenly comprised of dendrites from susceptible and resilient animals, was of similar size to the control set, and was randomly selected using a random number generator.

---

**Fig S1.**
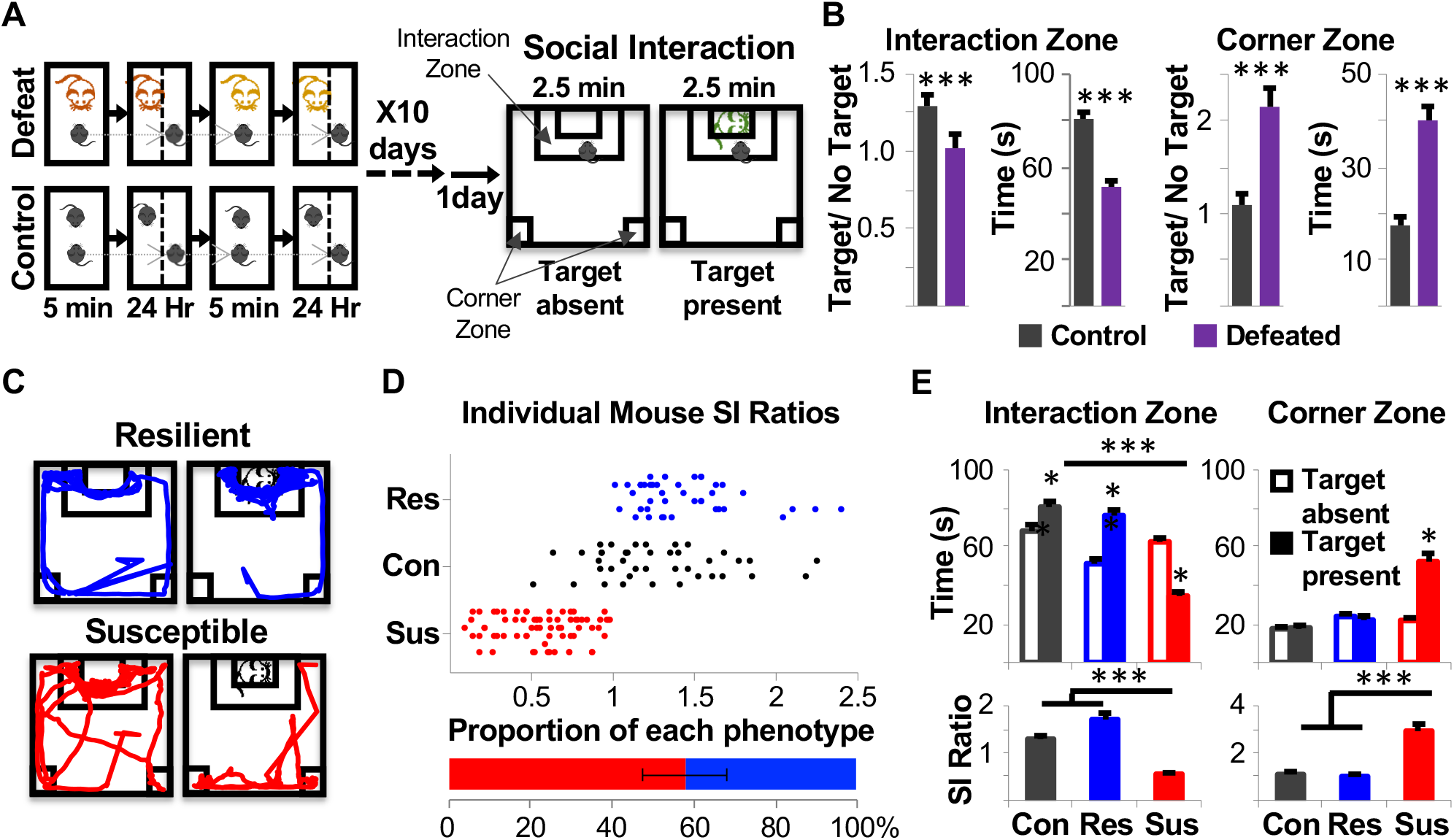
Overview of Chronic Social Defeat Stress. (A) Each day for 10 days an intruder C57 male mouse is placed in the cage of a novel aggressive CD1 retired breeder mouse for 5-10 minutes per day. The C57 is then moved to the other side of a perforated barrier which allows for sensory, but not physical, interaction between the C57 and CD1 for the remaining 24 hour day. After 10 days, the C57 is singly housed or re-group housed with prior cage-mates with *ad libitum* access to food and water for 24 hours before social interaction (SI) testing. For SI, the intruder C57 is first placed into an arena containing an empty wire-mesh enclosure and allowed to explore freely for 150 seconds. The intruder C57 is then removed from the arena and a non-aggressive CD “target” mouse is placed in the enclosure. The C57 is returned to the arena allowed another 150 seconds of exploration. (B) As a group, defeated mice show a pattern of social avoidance, with overall lower interaction time (two-tailed unpaired T-test, p<0.0001), lower interaction ratio (SI ratio, two-tailed unpaired T-test, p=0.0002), higher corner zone time (two-tailed unpaired T-test, p<0.0001), and higher corner ratio (two-way unpaired T-test, p<0.0001). Bar graphs show means of groups with SEM bars. (C) Susceptible mice are defined as mice with SI<1. Resilient mice are defined as mice SI ≥ 1. Example of track tracing of resilient and susceptible mice when target is absent versus present. Resilient (blue) mice spend more time in the interaction zone when target is present while susceptible mice (red) avoid the zone when a target mouse is present. (D) Distribution of SI ratios of individual control (black), resilient (blue), and susceptible (red) mice. Stressed mice are predominantly susceptible (58% ± 10% SEM). (E) Two-way ANOVAs were used to compare interaction and corner times when target was present versus absent and one-way ANOVAs were used to compare ratio scores. Bonferroni post-hoc tests were performed on all ANOVAs. Control mice are socially preferent and spend more time in interaction zone when a target mouse is present. By definition resilient mice spend more time in the interaction zone when a target mouse is present, while susceptible mice spend less time [two-way repeated measures ANOVA, GroupXTarget F(2,156)=124.6 p<0.001, group main effect F(2,156)=36.56 p<0.0001, target main effect F(1,156)=5.161 p=0.0245, C p=0.002, R p<0.0001, S p<0.0001]. SI ratios are lower for susceptible mice but do not differ from control for resilient mice [one-way ANOVA, F(2,156)=20.86 p<0.0001, post-hoc test: C vs S p=0.0015, R vs S p<0.0001, C vs R p= 0.3935]. Conversely, susceptible mice spend more time in corner zones [two-way repeated measures ANOVA, GroupXTarget F(2,156)=33.97 p<0.001, group main effect F(2,156)=24.97 p<0.0001, target main effect F(1,156)=21.05 p<0.0001, C p=n.s., R p=n.s., S p<0.0001] and have higher corner time ratios [one-way ANOVA, F(2,156)=21.05 p<0.0001, post-hoc Bonferroni test: C vs S p<0.0001, R vs S p<0.0001, C vs R p= 0.83] compared to control and resilient mice.

**Fig S2.**
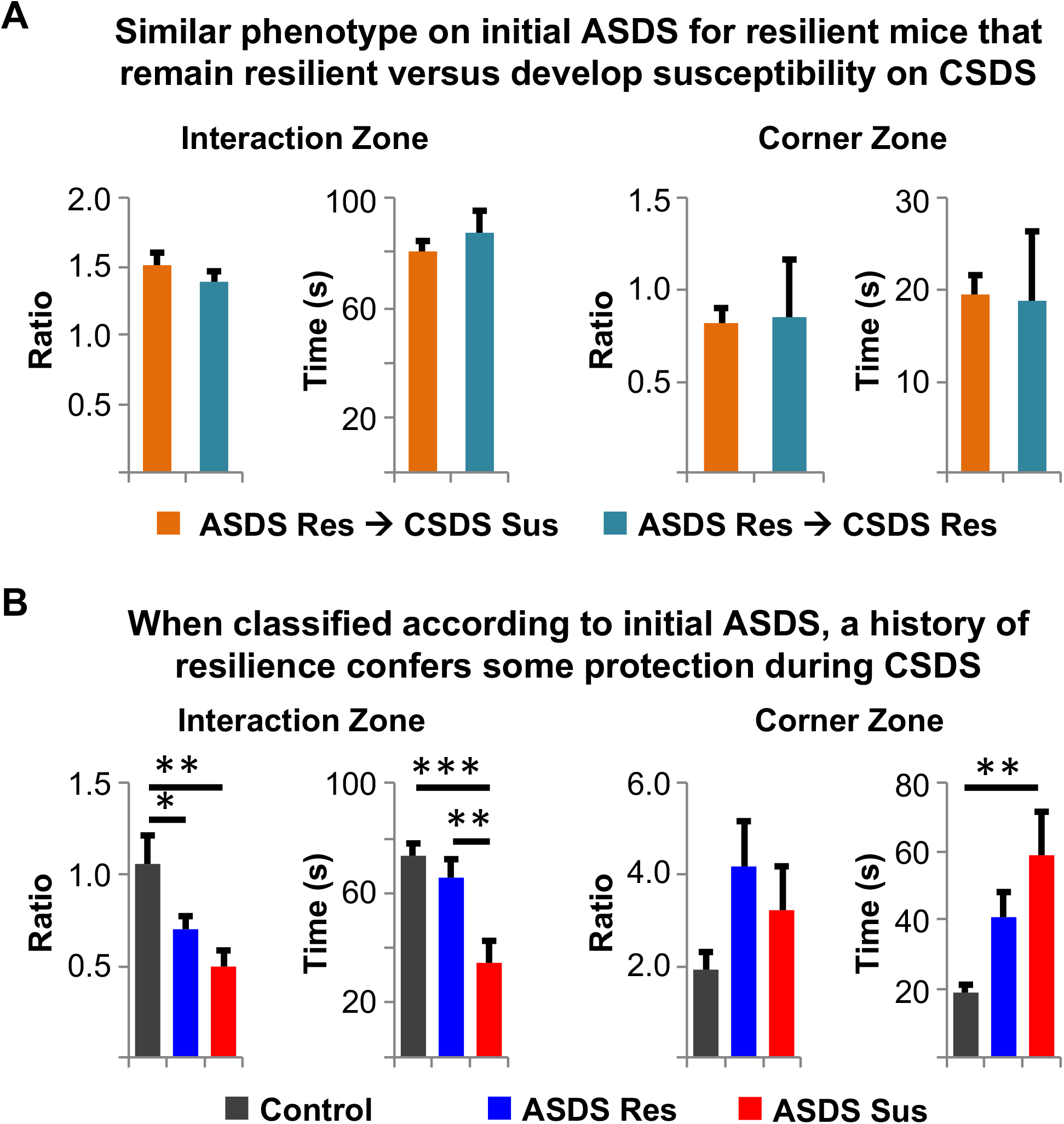
Comparison of behavioral phenotype on initial ASDS versus subsequent CSDS. C57 mice underwent ASDS and SI, allowed a 1 week recovery period, then placed in CSDS for 10 days and tested on SI on day 11. (A) Mice that were resilient after ASDS and remained resilient after CSDS (teal) had no significant differences in SI behavior after ASDS compared to mice that were resilient after ASDS, but became susceptible after CSDS (orange; two-tailed unpaired T-test, SI ratio p=0.55, interaction zone duration p=0.46, corner duration p=0.9, corner ratio p=0.89). (B) Mice that were identified as resilient after ASDS where somewhat protected from developing severe social avoidance following CSDS compared to mice classified as susceptible after ASDS [one-way ANOVA, SI ratio: F(2,61)=8.847 p=0.0004, post-hoc Bonferroni test: C vs S p=0.006, R vs S p=0.544, C vs R p= 0.0011, interaction duration: F(2,61)=8.106 p=0.0007, post-hoc Bonferroni test: C vs S p<0.0001, R vs S p=0.056, C vs R p= 0.022, corner ratio: F(2,60)=6.2.54 p=0.0872, post-hoc Bonferroni test: C vs S p=0.1755, R vs S p=0.3347, C vs R p= 0.039, corner duration: F(2,60)=5.245 p=0.008, post-hoc Bonferroni test: C vs S p=0.0011, R vs S p=0.4176, C vs R p= 0.0083]. Bar graphs show means of groups with SEM. C=control, R=resilient, S=susceptible While “super-resilient” mice cannot be predicted on the basis of individual behavioral indices, a classifier combining multiple scores from the initial ASDS can be implemented. First, animals with freezing rather than avoidant behavior are excluded. Resilience is then defined to meet all three of the following criteria: SI ≥ 1, absolute time interacting with novel target mouse ≥ 60 sec, and absolute time spent in corners when novel target mouse present < 30 sec (see Extended Methods for further details). This classifier identifies “super-resilient” mice with 83% sensitivity and 59% specificity.

**Fig S3.**
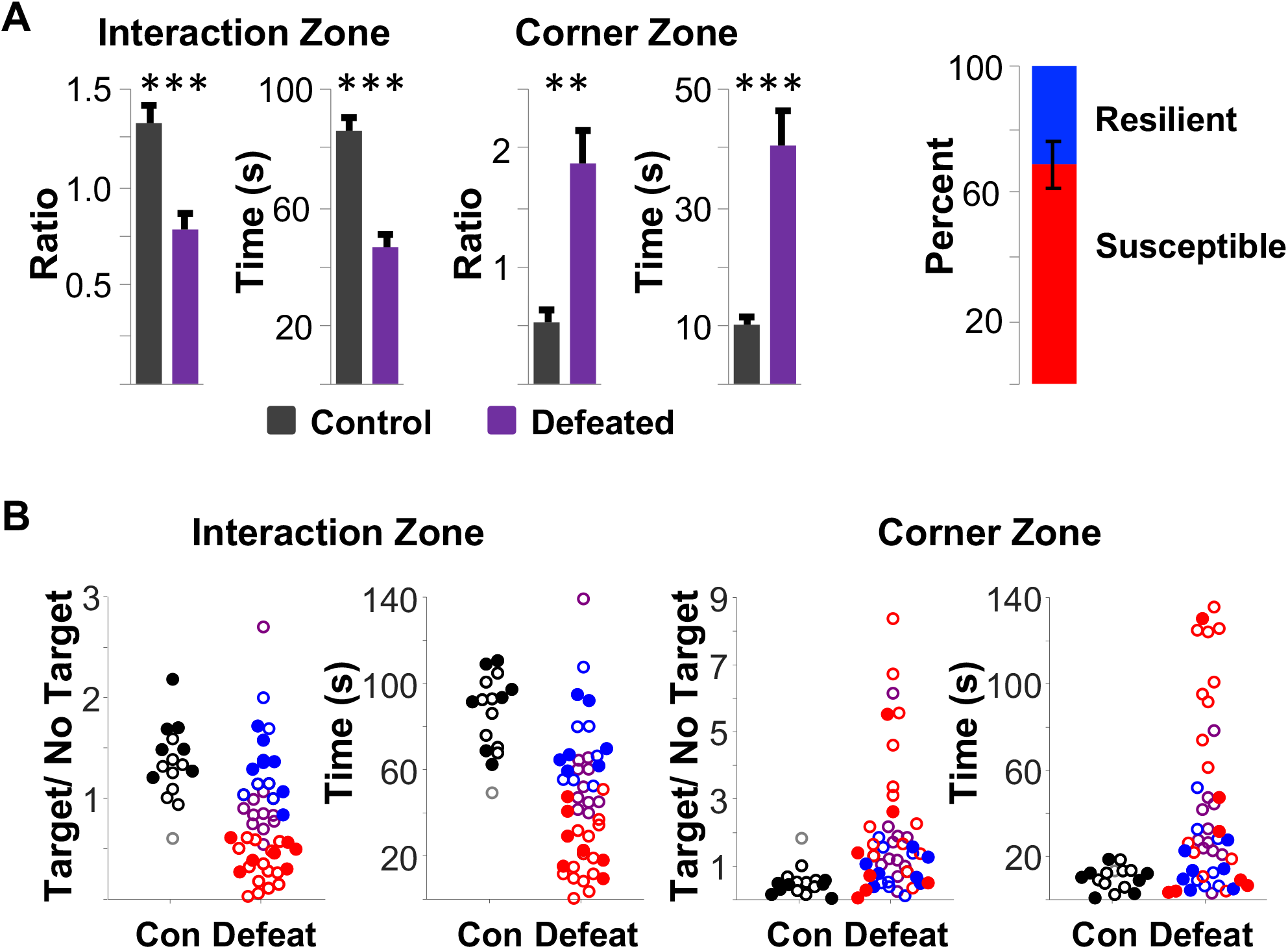
Stereotactic BLA injections for viral-mediated tract tracing do not have an effect on ASDS. (A) Times and ratios spent in interaction and corner zones for mice in viral-mediated tract-tracing experiment (N=37). Pattern of social avoidance and percent resilience (defined as SI>1, 68.95% ± 7.39) is similar to the overall distribution of mice undergoing ASDS (Figure 1 in main text) (two-tailed unpaired t-test, interaction duration: p<0.0001, corner duration: p<0.0001, SI ratio: p<0.0001, corner ratio: p=0.006) (B) The distribution of behavioral responses of mice in viral-mediated tract-tracing experiment. Closed circles indicate mice that met the multidimensional behavioral classifier and had confirmed BLA injection sites (n=7 control, n=7 resilient, n=7 susceptible). Open circles indicate mice that did not meet either behavioral and/or histological criteria. Black=control, blue=resilient, red=susceptible, gray=animal that displays freezing rather than avoidant behavior, defined as average velocity<1.8 cm/s.

**Fig S4.**
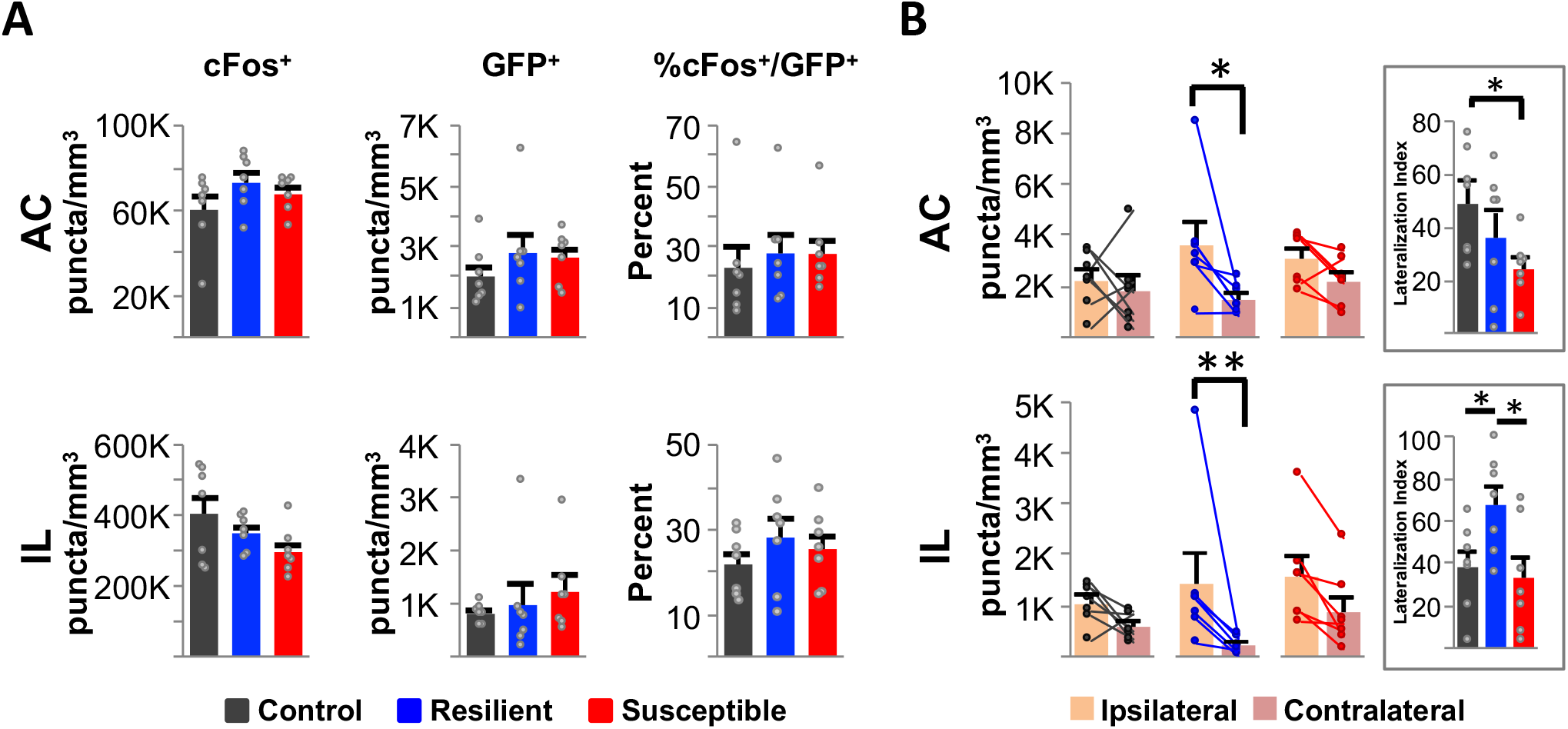
No differences in the activation of anterior cingulate (AC) or infralimbic cortex (IL) but trend of higher lateralization of cortical BLA-projecting neurons in resilient mice extends to these regions. (A) Expression of cFos, GFP, and cFos/GFP colocalization in AC and IL. There were no significant differences in GFP or cFos expression, or in cFos/GFP colocalization between control (C, black), resilient (R, blue), or susceptible (S, red) mice [one-way ANOVA, AC cFos+: F(2,18)=1.585 p=0.232, AC GFP+: F(2,18)=0.832 p=0.451, AC %cFos+/GFP+: F(2,18)=0.185 p=0.832, IL cFos+: F(2,18)=2.5118 p=0.1091, IL GFP+: F(2,18)=0.1344 p=0.8751, IL %cFos+/GFP+: F(2,18)=0.7172 p=0.5015]. Bar graphs show the mean puncta density per group with SEM error bars. Scatter plots show individual animal values. (B) Lateralization of GFP expression. Resilient mice had greater density of AC → BLA neurons in ipsilateral compared to contralateral AC [two-way repeated measures ANOVA, GroupXHemisphere F(2,18)=1.797 p=0.1943, group main effect F(2,18)=0.7479 p=0.4874, hemisphere main effect F(1,18)=8.875 p=0.008, R p=0.0147, S p=0.57, C p=0.99]. Resilient mice also had greater density of IL → BLA neurons in ipsilateral compared to contralateral IL [two-way repeated measures ANOVA, GroupXHemisphere F(2,18)=1.202 p=0.3237, group main effect F(2,18)=0.8317 p=0.4514, hemisphere main effect F(1,18)=14.54 p=0.0013, R p=0.0096, S p=0.1967, C p=0.6864]. Inset: Lateralization of hemispheres was calculated as density of mPFC → BLA neurons in ipsilateral hemisphere minus density of mPFC → BLA neurons in the contralateral hemisphere divided by combined density in both hemispheres. Susceptible mice had significantly lower lateralization than control mice in AC → BLA neurons [one-way ANOVA, F(2,18)=2.81 p=0.087, C v S p=0.015]. Resilient mice had significantly higher lateralization than control and susceptible mice in IL → BLA [one-way ANOVA, F(2,18)=4.2975 p=0.0298, C v R p=0.0291, R v S p=0.0248). Bar graphs show the mean puncta density per group with SEM error bars. Scatter plots show individual paired hemisphere values. Taken together, observed higher lateralization of PL→BLA neurons from resilient versus susceptible mice generalizes to the rest of the mPFC.

**Fig S5.**
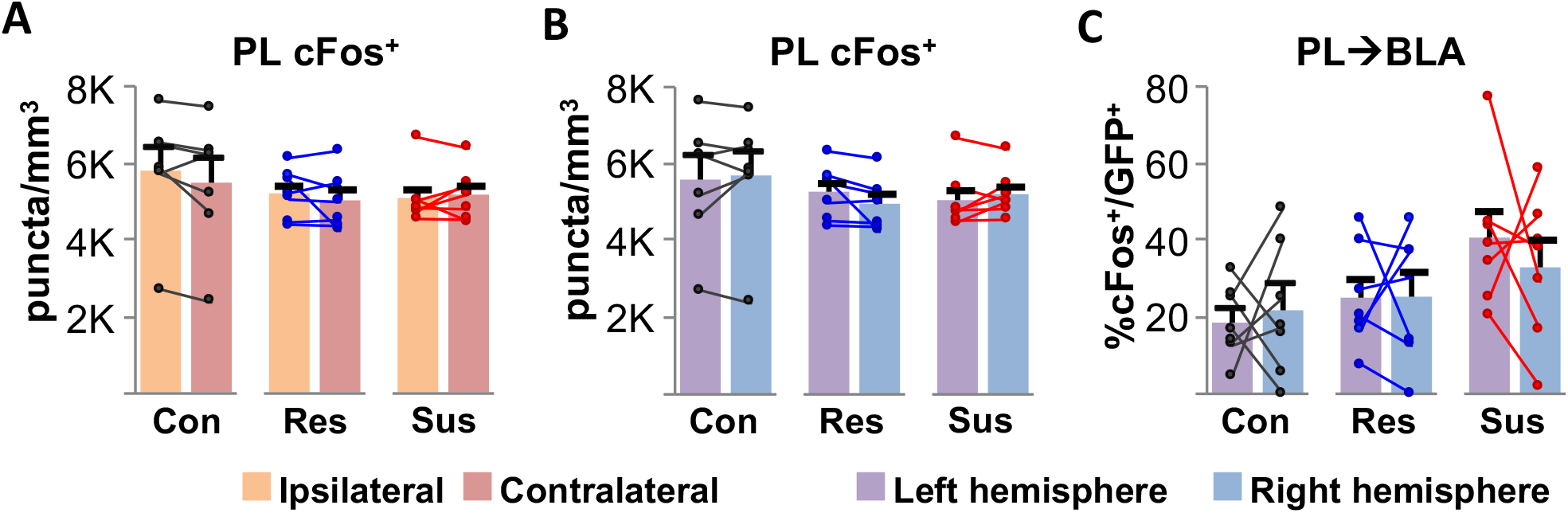
Observed PL→BLA ipsi-versus contra-lateral activation differences are not mediated by overall differences in PL ipsi-/contra-lateral hemispheric activation or by left/right hemispheric activation differences. (A) There was no significant difference in cFos puncta density between ipsilateral and contralateral hemispheres of control, resilient, or susceptible mice [two-way repeated measure ANOVA, GroupXHemisphere F(2,18)=1.186 p=0.3281, group main effect F(2,18)=0.5748 p=0.5728, hemisphere main effect F(1,18)=1.581 p=0.2247]. (B) There was no significant difference in cFos puncta density between left and right hemisphere of control, resilient, or susceptible mice [two-way repeated measures ANOVA, GroupXHemisphere F(2,18)=1.901 p=0.1782, group main effect F(2,18)=0.5748 p=0.5728, hemisphere main effect F(1,18)=0.023 p=0.881]. (C) There was no significant difference in percentage of GFP puncta colocalized with cFos puncta in the left or right hemisphere, although consistant with the higher proportion activation of PL→BLA neurons from susceptible described in the main text, there was a main group effect [two-way repeated measures ANOVA, GroupXHemisphere F(2,18)=0.4323p=0.6556, group main effect F(2,18)=3.98 p=0.037, hemisphere main effect F(1,18)=0.0818 p=0.7782]. Bar graphs show the mean puncta density per group with SEM error bars. Scatter plots show individual paired hemisphere values.

**Fig S6.**
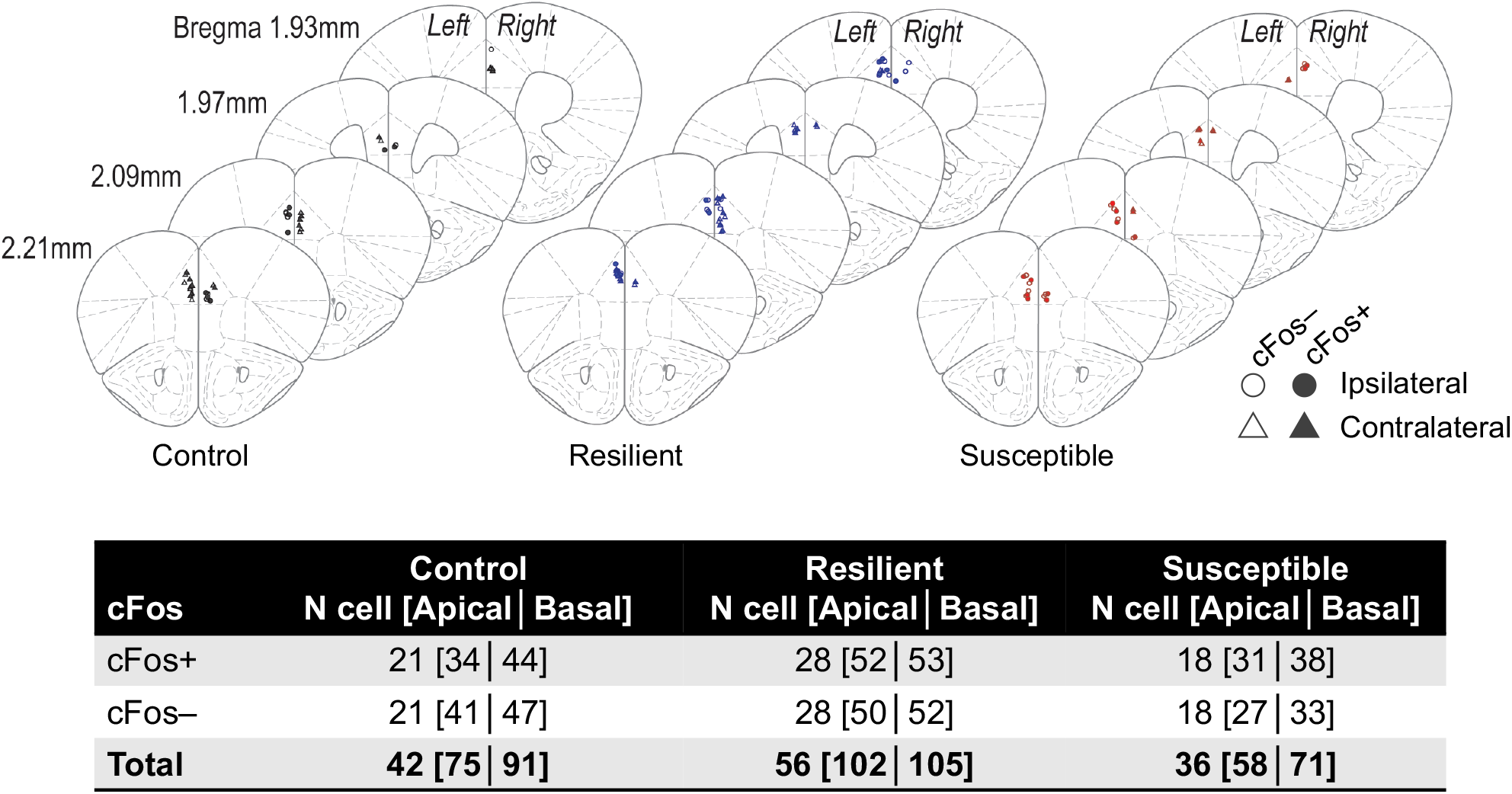
Location and number of PL→BLA neurons targeted for dendritic spine analysis. Plates were taken from Paxinos atlas to demonstrate locations of cells included in dendritic analyses. Circles indicate that the cell was ipsilateral to the viral injection site while triangles indicate that the cell was located in the contralateral hemisphere from the injection site. Filled circles or triangles indicate cFos immunoreactive (cFos+) neuron while empty shapes indicate cFos immunoreactive (cFos−) neuron. A total of 502 dendrites from 156 neurons with somas located in layers II-V were targeted for high resolution imaging. Of these, 20 of the control neurons, 25 of the resilient neurons, and all susceptible neurons are location paired “neighbor” neurons to allow for paired analyses for the evaluation of activity-dependent changes in spines.

**Fig S7.**
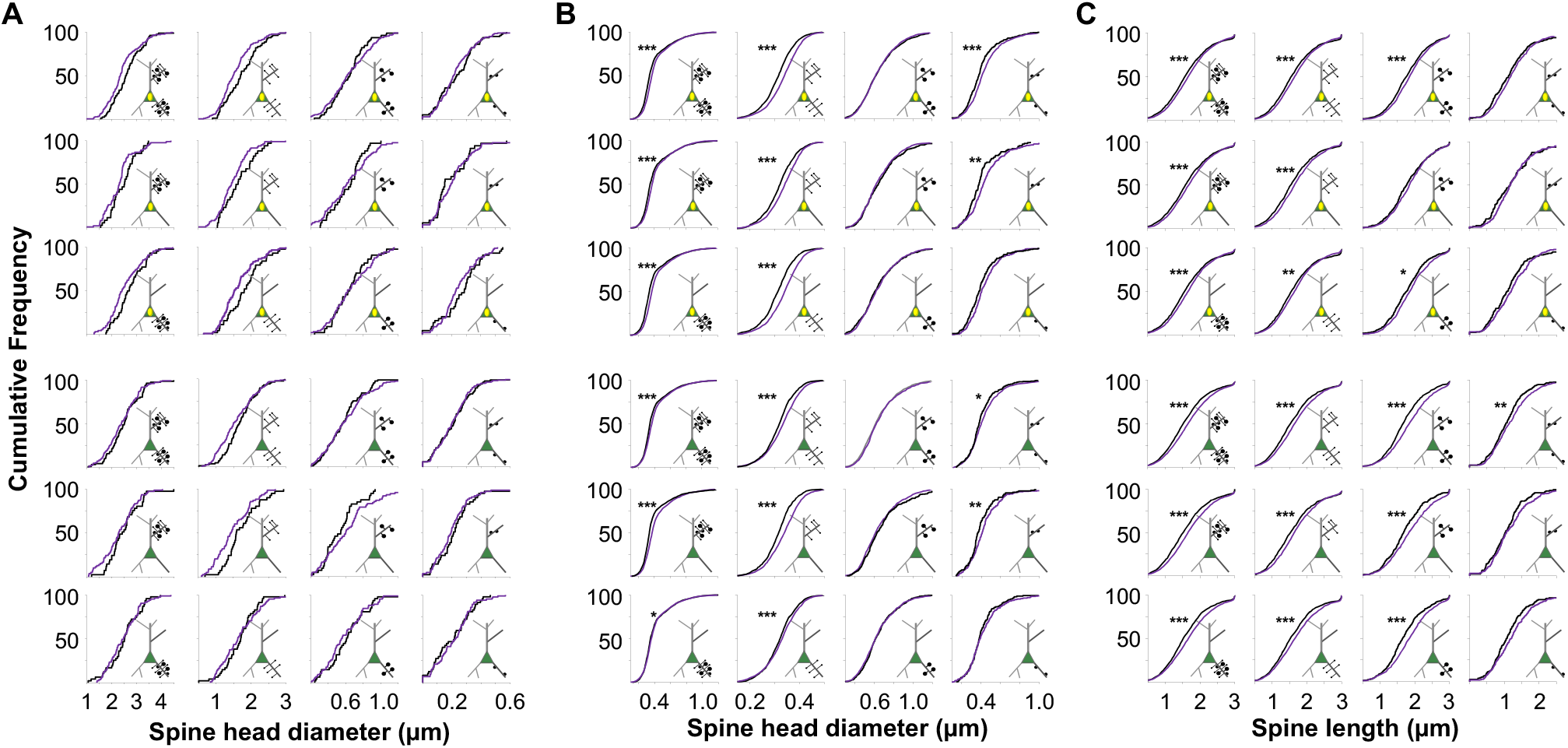
Dendritic spine density and morphology comparison of Control vs Stress. Cartoon neuron insets indicate cFos+ (yellow oval in the center of green s) or cFos− (no yellow oval in green triangle), spine subtype (thin= long and small, mushroom= long and large, stubby= short and small) and apical (spines on upper dendrites) or basal (spines on lower dendrites). All reported p-values represent Kolmogorov-Smirnov (K-S) tests with Bonferroni correction. (A) Cumulative distribution frequency (CDF) plots comparing spine densities of dendrites from control (black) versus combined stress group (purple, susceptible+resilient mice). There were no significant differences in dendritic spine density between stress and control mice in any spine subtype on apical or basal dendrites of cFos+ or cFos− neurons. (B) CDF plots comparing spine head diameters between control and stress. Stress group mice had larger head diameters of thin spines on apical and basal dendrites of cFos+ and cFos− neurons (cFos+ apical/basal all subtypes p<0.0001, cFos+ apical all p<0.0001, cFos+ basal all p<0.0001, cFos+ apical/basal thin p<0.0001, cFos+ apical thin p<0.0001, cFos+ basal thin p<0.0001, cFos− apical/basal all p<0.0001, cFos− apical all p<0.0001, cFos− basal all p=0.0338, cFos− apical/basal thin p<0.0001, cFos− apical thin p<0.0001, cFos− basal thin p<0.0001). Stress group mice also had greater spine head diameter of stubby spines on combined apical and basal dendrites of cFos+ and cFos− neurons and on apical dendrites of cFos+ and cFos− neurons (cFos+ apical/basal stubby p<0.0001, cFos+ apical stubby p=0.0038, cFos− apical/basal stubby p=0.024, cFos− apical stubby p=0.010). As reported in the main text, there was no significant difference in spine head diameter of mushroom spines. (C) CDF plots comparing spine lengths between control and stress. Stress group mice had longer apical and basal thin spines on cFos+ and cFos− neurons (cFos+ apical/basal all p<0.0001, cFos+ apical all p<0.0001, cFos+ basal all p<0.0001, cFos+ apical/basal thin p<0.0001, cFos+ apical thin p<0.0001, cFos+ basal thin p=0.006, cFos− apical/basal all p<0.0001, cFos− apical all p<0.0001, cFos− basal all p<0.0001, cFos− apical/basal thin p<0.0001, cFos− apical thin p<0.0001, cFos− basal thin p<0.0001). Stress group mice had longer mushroom spines on basal dendrites of cFos+ neurons, and apical and basal dendrites of cFos− neurons (cFos+ apical/basal mushroom p<0.0001, cFos+ basal mushroom p=0.032, cFos− apical/basal mushroom p<0.0001, cFos− apical mushroom p<0.0001, cFos− basal mushroom p<0.0001). Stress group mice had longer stubby spines on cFos− neurons when apical and basal dendrites were combined (cFos− apical/basal stubby p=0.035).

**Fig S8.**
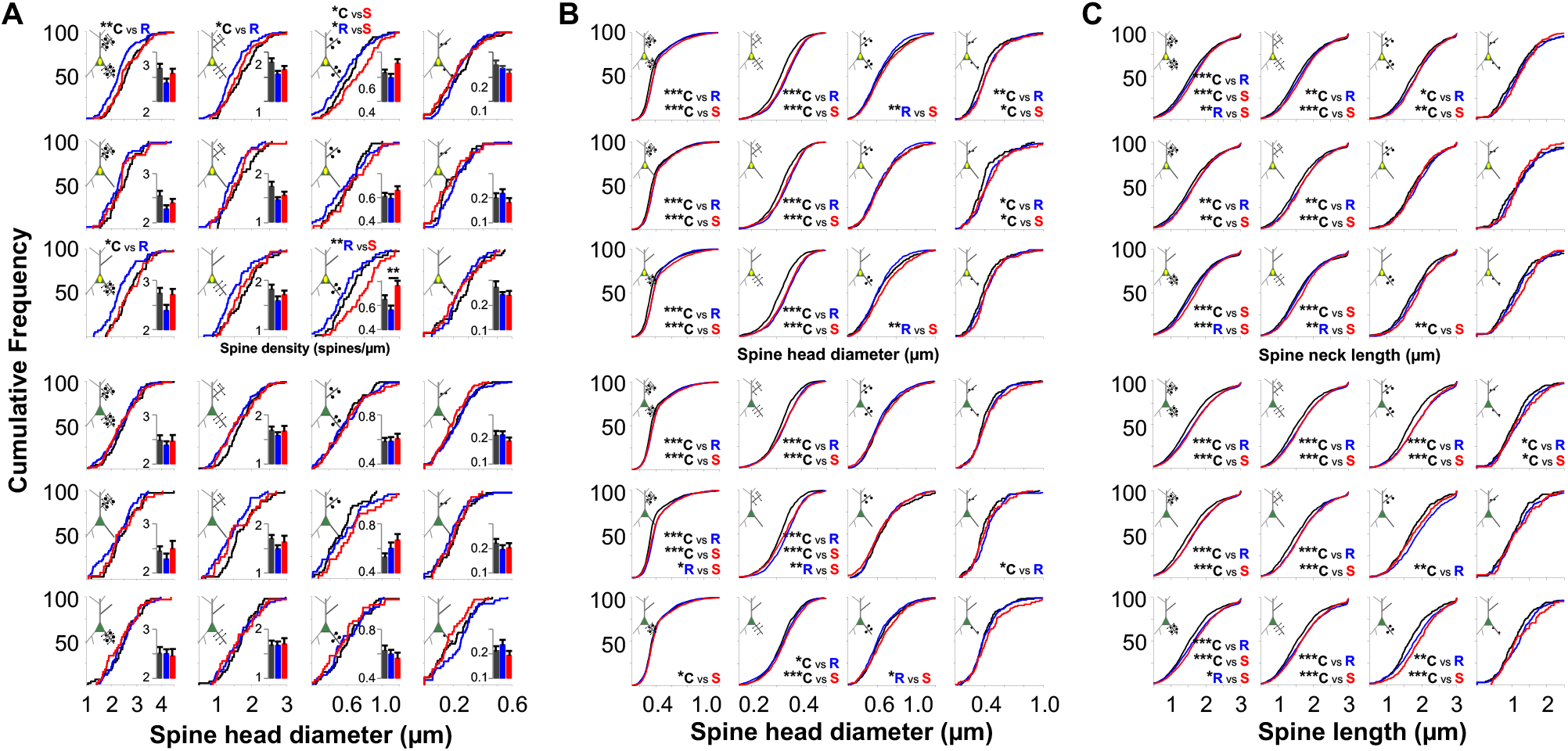
Dendritic spine density and morphology differences with all spines included. Expanded **Figure 5** to include CDF plots of combined all subtype spines on apical and basal dendrites (left column), combined all thin, mushroom, and study spines on both apical and basal dendrites (top row and forth row), and all spines combined (left upper corner graphs in each panel). All reported p-values represent K-S tests with Bonferroni correction. (A) CDF plots of spine densities. In addition to the findings reported in the main text, cFos positive neurons from resilient (R) mice had lower total spine densities (p=0.0027), lower basal combined spine densities (p=0.045), and lower thin combined spine densities (p=0.012) than control (C). Susceptible (S) mice had higher combined apical and basal mushroom spine density than control (p= 0.012) and resilient mice (p=0.012) on cFos+ neurons. cFos– continued to not display any dendritic spine differences. Bar graphs display group means with SEM error bars and no additional findings were identified than those reported in main text. (B) CDF plots of spine head diameter. Resilient and susceptible mice had larger head diameter of total combined spines, combined thin spines, combined stubby spines, all subtypes on basal dendrites, and all subtypes on apical dendrites of cFos+ neurons (cFos+ basal/apical all C v R p<0.001, C v S p<0.0001, cFos+ basal/apical thin C v R p<0.0001, C v S p<0.0001, cFos+ basal/apical stubby C v R p=0.0033, C v S p=0.018, cFos+ apical all C v R p<0.0001, C v S p<0.0001, cFos+ basal all C v R p<0.0001, C v S p<0.0001). Susceptible mice also had higher spine head diameter of combined mushroom spines on cFos+ neurons compared to resilient (p=0.0064). Controls had smaller head diameter of total combined spines, combined thin spines, and combine subtypes on apical dendrites of cFos− neurons compared to resilient and susceptible (cFos− basal/apical all C v R p<0.0001, C v S p<0.0001, cFos− basal/apical thin C v R p<0.0001, C v S p<0.0001, cFos− apical all C v R p<0.0001, C v S p<0.0001). Control mice also had smaller head diameter of combined subtypes on basal dendrites of cFos− neurons compared to susceptible (p=0.032). (C) CDF plots of spine lengths. Resilient and susceptible mice had longer spine length of total combined spines, combined thin spines, combined mushroom spines, all subtypes on basal dendrites, and all subtypes on apical dendrites of cFos+ and cFos− neurons compared to control mice (cFos+ basal/apical all C v R p<0.001, C v S p<0.0001, cFos+ basal/apical thin C v R p=0.0018, C v S p<0.0001, cFos+ basal/apical mushroom C v R p=0.0023, C v S p=0.0063, cFos+ apical all C v R p=0.002, C v S p<0.0001, cFos+ basal all C v S p<0.0001, cFos− basal/apical all C v R p<0.0001, C v S p<0.0001, cFos− basal/apical thin C v R p<0.0001, C v S p<0.0001, cFos− basal/apical mushroom C v R p<0.0001, C v S p<0.0001, cFos− basal/apical stubby C v R p=0.037, C v S p=0.036, cFos− apical all C v R p<0.0001, C v S p<0.0001, cFos− basal all C v R p<0.0001, C v S p<0.0001). Susceptible and resilient mice also had longer spine length than control mice of combined stubby spines on cFos− neurons (C v R p<0.0001, C v S p<0.0001). Susceptible had significantly longer spine necks of combined subtypes on basal dendrites compared to resilient (K-S test with Bonferroni correction, p=0.033).

**Fig S9.**
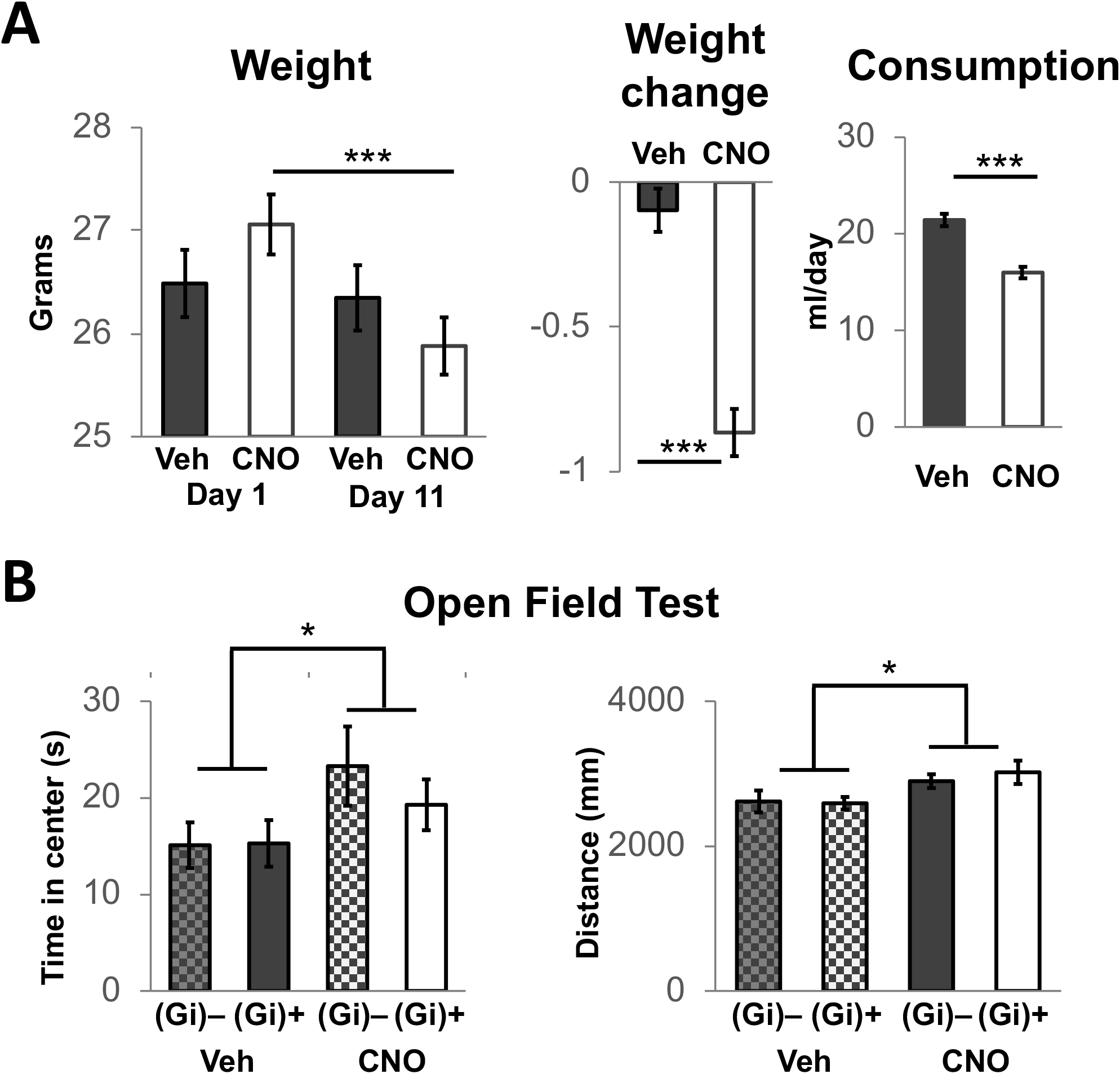
Off-target effects of chronic CNO. (A) Effects of CNO on weight. Mice that received CNO in their drinking water (n=40) had significantly lower weights (two-way repeated measures ANOVA, TimeXDrug F(1,76)=46.92 p<0.0001, drug main effect: F(1,76)=0.0157 p=0.9006, time main effect: F(1,76)=75.45 p<0.0001, CNO Day 11 v Day 1 p<0.0001) and a weight loss (two-tailed unpaired T-test, p<0.0001) at the end of 10 day administration compared to mice that only received vehicle (aspartame) in their drinking water (N=38). CNO treated mice also consumed less fluid (two-tailed unpaired T-test, p<0.0001). (B) Open field testing revealed a significant DREADD-independent drug effect of increased time spent in the center area [10×10cm, two-way ANOVA, DREADDXDrug F(1,76)=0.5054 p=0.4794, DREADD main effect: F(1,76)=0.4205 p=0.5187, drug main effect: F(1,76)=4.275 p=0.0422] and increased locomotor function [two-way ANOVA, DEADDXDrug F(1,76)=0.2596 p=0.6119, DREADD main effect: F(1,76)=0.1128 p=0.738, drug main effect: F(1,76)=6.027 p=0.0165) and spent less time in the center of the arena] in chronically CNO treated mice.

## Notes

### Competing Interest Statement

The authors have declared no competing interest.

